# Uranium-bearing dust induces differentiation and expansion of enteroendocrine cells in human colonoids

**DOI:** 10.1101/2023.08.10.552796

**Authors:** Roger Atanga, Lidia L. Appell, Fredine T. Lauer, Adrian Brearley, Matthew J. Campen, Eliseo F. Castillo, Julie G. In

## Abstract

Chronic exposure to environmental toxins and heavy metals has been associated with intestinal inflammation, increased susceptibility to pathogen-induced diseases, and higher incidences of colorectal cancer, all of which have been steadily increasing in prevalence for the past 40 years. The negative effects of heavy metals on barrier permeability and inhibition of intestinal epithelial healing have been described; however, transcriptomic changes within the intestinal epithelial cells and impacts on lineage differentiation are largely unknown. Uranium exposure remains an important environmental legacy and physiological health concern, with hundreds of abandoned uranium mines located in the Southwestern United States largely impacting underserved indigenous communities. Herein, using human colonoids, we defined the molecular and cellular changes that occur in response to uranium bearing dust (UBD) exposure. We used single cell RNA sequencing to define the molecular changes that occur to specific identities of colonic epithelial cells. We demonstrate that this environmental toxicant disrupts proliferation and induces hyperplastic differentiation of secretory lineage cells, particularly enteroendocrine cells (EEC). EECs respond to UBD exposure with increased differentiation into *de novo* EEC sub-types not found in control colonoids. This UBD-induced EEC differentiation does not occur via canonical transcription factors *NEUROG3* or *NEUROD1.* These findings highlight the significance of crypts-based proliferative cells and secretory cell differentiation as major colonic responses to heavy metal-induced injury.

## INTRODUCTION

Uranium mining in the United States arose in the 1940s, peaked in the 1970s, and subsided in the late 1980s, with extensive activity concentrated in Navajo, Puebloan, and other indigenous lands in the Southwest (Brugge and Goble, 2002). To date, over 500 mines in this region are considered abandoned uranium mines (AUM). Heavy metals from AUMs transfer into the sediment and groundwater (Eadie and Kaufmann, 1977; Lin et al., 2020), with non-fissile uranium (^238^U) levels in surrounding communities exceeding maximum contamination levels (30 μg/L) dictated by the US Environmental Protection Agency. For example, well water in Nambe Pueblo (Santa Fe, New Mexico) once contained 1.2 mg/L ^238^U (Hakonson-Hayes et al., 2002). Even partially remediated uranium mines degrade over time and cause heavy metal contamination of the surrounding sediment and groundwater (Schaider et al., 2014). This has led to unacceptable ^238^U levels found in community water sources throughout the US, even in aquifers that are not near uranium mines (Nolan and Weber, 2015; Ravalli et al., 2022).

Although ^238^U is weakly radioactive, it primarily negatively influences human health as a chemotoxicant and can accumulate in various organs (Erdei et al., 2019; Rump et al., 2019; WHO, 2001). The communities that reside near AUMs are chronically exposed to particulate dust with varying levels of uranium (referred to as uranium-bearing dust, UBD) via inhalation and ingestion (Ma et al., 2020; Zychowski et al., 2018), increasing their overall health risks. Epidemiological and translational studies in rodents found that chronic environmental exposure to UBD causes: impaired neurological function (Legrand et al., 2016), pulmonary damage (Monleau et al., 2006), hepatotoxicity (Souidi et al., 2005), renal damage (Zamora et al., 2009), and reproductive defects (Wang et al., 2020a).

Specific to the gut, chronic exposure to environmental toxins and heavy metals has been linked to intestinal inflammation (Legaki and Gazouli, 2016), increased susceptibility to pathogen-induced diseases (Chiu et al., 2020; Wales and Davies, 2015), circulating inflammatory potential (Harmon et al., 2017), and colorectal cancer (Wagner et al., 2011). Additionally, heavy metal exposure has been shown to negatively impact the microbiota (Richardson et al., 2018), immune cells (Dublineau et al., 2007; Medina et al., 2020), and the intestinal epithelial barrier (Bolan et al., 2021). Yet, the molecular changes that lead to these pathologies are undefined. ^238^U, via UBD, enters the intestine via ingestion or inhalation. Inhaled sedimentary particles are transported to the epiglottis by tracheobronchial cilia, then swallowed (Entwistle et al., 2019). Reported amounts and rate of uranium absorption in the digestive tract are fairly low, from 0.1 – 7% (Traber et al., 2014), bringing into question how systemic health impacts arise from direct toxicity to the gut. In mice, ingested uranium was primarily absorbed in the lower intestinal tract (colon) (Medina et al., 2020), suggesting ^238^U-related cellular changes may originate in the colon.

The intestine is the largest immune organ in humans (Furness et al., 2013) and directly communicates with the brain via hormone-secreting enteroendocrine cells and the enteric nervous system (Kuwahara et al., 2020; Latorre et al., 2016). Thus, detrimental effects to the intestine can indirectly affect systemic health. The UBD utilized in this study is toxicologically relevant (**Supplemental Figure 1**) as it was obtained from a structure in Paguate, NM, a village in the Laguna Pueblo located next to one of the largest open-pit uranium mines in the United States, the Jackpile Mine (Moore-Nall, 2015).

Human colonoid cultures are a tractable, epithelial-only model that can indefinitely proliferate due to the presence of adult intestinal stem cells (Sato et al., 2011), making them a valuable model to study how the epithelium responds to intestinal injury and the factors involved in epithelial regeneration. The current lack of understanding on how heavy metals, specifically in particulate form like UBD, affect intestinal epithelia suggest a novel approach is needed. We use human colonoids as a relevant model of human intestine to characterize UBD-induced transcriptomic changes and show that human colonoids are a relevant pathophysiological model for studies in host-toxicant interactions.

Enteroendocrine cells (EECs) are rare hormone-producing cells in the gut (Osinski et al., 2022; Tsang et al., 2022). They are distributed throughout the intestinal epithelium, making up only approximately 1% of the epithelial cell population. Thus, EEC function is difficult to study in whole animals. Recent studies showed that intestinal organoids are a physiologically relevant model to study EEC differentiation and function (Bellono et al., 2017; Goldspink et al., 2018a). Intestinal organoids produce the various EEC sub-types, retain regional specificity to the gut hormones produced, and respond to a variety of physiological stimuli (Basak et al., 2017).

In this study, we used single cell RNA-sequencing (scRNA-seq) to characterize UBD-induced changes to intestinal epithelial cell lineages in adult stem cell-derived human colonoids. From scRNA-seq analysis and validation with RNA fluorescence in situ hybridization and immunostaining, we found that UBD disrupts homeostasis by depleting proliferative cells and inducing differentiation towards secretory lineages. Colonic EECs expand and further differentiate to sub-types that are not present in control colonoids. This *de novo* enteroendocrine expansion is independent of the canonical transcription factors, *NEUROD1* and *NEUROG3*.

## RESULTS

### Acute UBD exposure is not cytotoxic to human colonoids

To determine if UBD induces cell death or morphological changes in human colonoids, three biologically distinct human colonoid cultures were treated with vehicle (control) or 50 µg/ml UBD (resuspended in organoid expansion media) overnight (approximately 18 h). Brightfield imaging of both conditions showed no changes in gross morphology after acute UBD exposure (**Fig. 1A**). Propidium iodide (PI) staining was used to visualize dead cells. No significant difference in PI-positive cells was observed between control and UBD treated colonoids (**Fig. 1B**).

**Fig 1.**
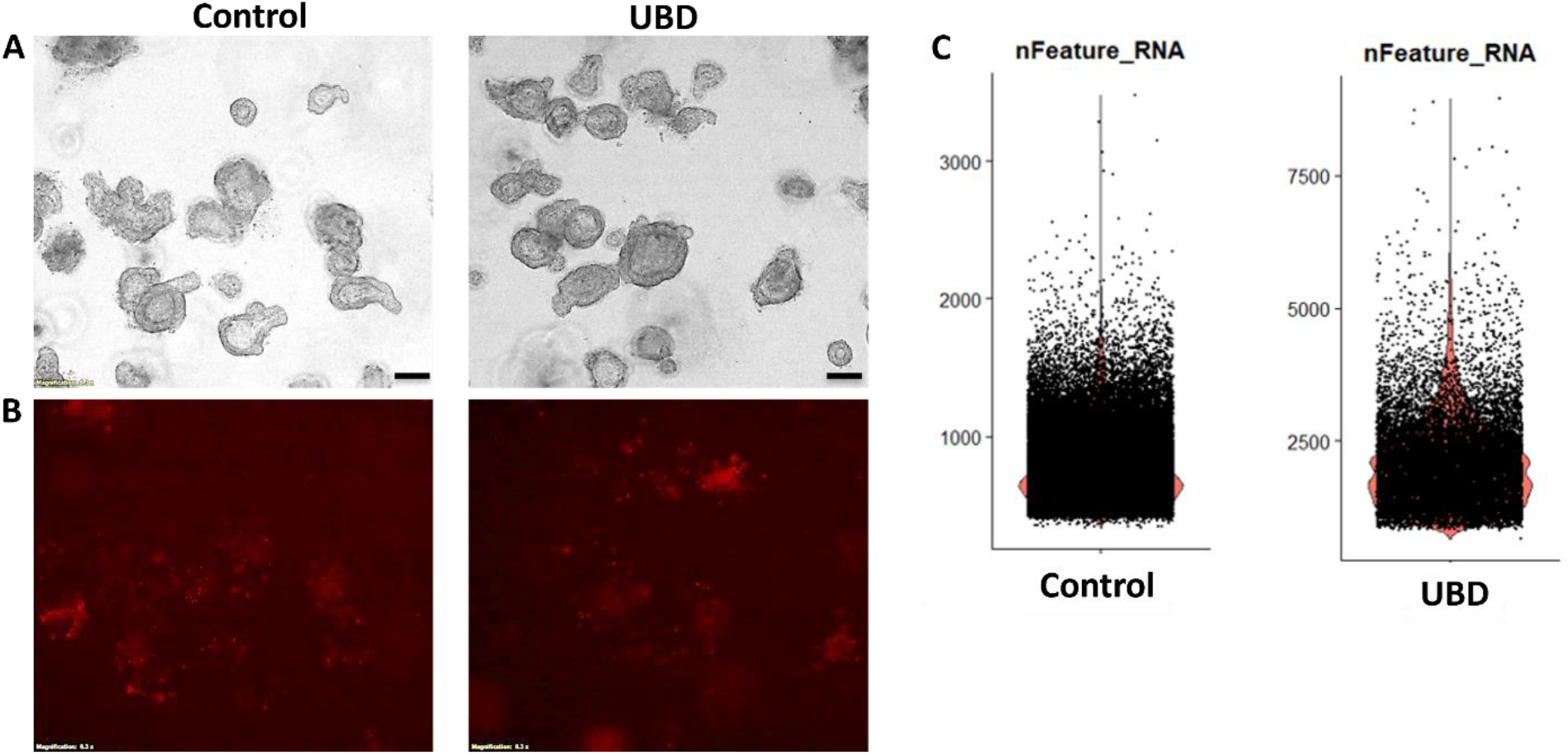
Acute UBD exposure does not increase cell death in human colonoids but increases gene expression per cell. **A)** and **B)** Human colonoids were treated with vehicle (control) or UBD overnight; scale bar = 200 µm, N ≥ 3. **A)** Representative brightfield images show no gross differences between control and UBD exposed colonoids. **B)** The colonoids were stained with propidium iodide (dead cells). Representative propidium iodide staining (red) shows no significant differences in cell death between the two conditions. **C)** Representative violin plots for gene expression numbers per cell shows control colonoids express <3000 while some UBD-exposed express >5000.

To understand if UBD is changing specific cell lineages, we performed droplet based single cell RNA-sequencing (scRNA-seq) on three biologically unique colonoid lines (**Supplemental Table 1)** under control or UBD treated conditions. Despite the lack of changes in cell death and overall morphology, computationally there was a stark difference in cellular gene expression. Cells in control colonoids expressed <3000 genes while some UBD treated cells expressed >5000 genes (**Fig. 1C**). These observations indicate that while acute UBD exposure has no cytotoxic effects on human colonoids, it induces changes in gene expression in colonic epithelial cells.

### Acute UBD exposure disrupts homeostasis and induces differentiation

Observing the 2-fold increase in gene expression per cell after UBD exposure, we sought to investigate the molecular and transcriptomic changes that occur following UBD treatment. Graph-based clustering using the Louvain algorithm and marker-gene analysis revealed few clusters in control colonoids that primarily represented cells founds in the deep crypt, as shown in the representative control colonoid UMAP (**Fig. 2A, left**). Clusters were manually identified using marker genes and published datasets (Beumer et al., 2020; Burclaff et al., 2022; Wang et al., 2020b) as: cycling transit amplifying (TA) cells, stem cells, and two clusters of secretory progenitor cells (**Dataset 1**). The colonoids represent non-differentiated epithelia since they were maintained in organoid expansion media which contains WNT3A/Rspo-1/Noggin.

**Fig 2.**
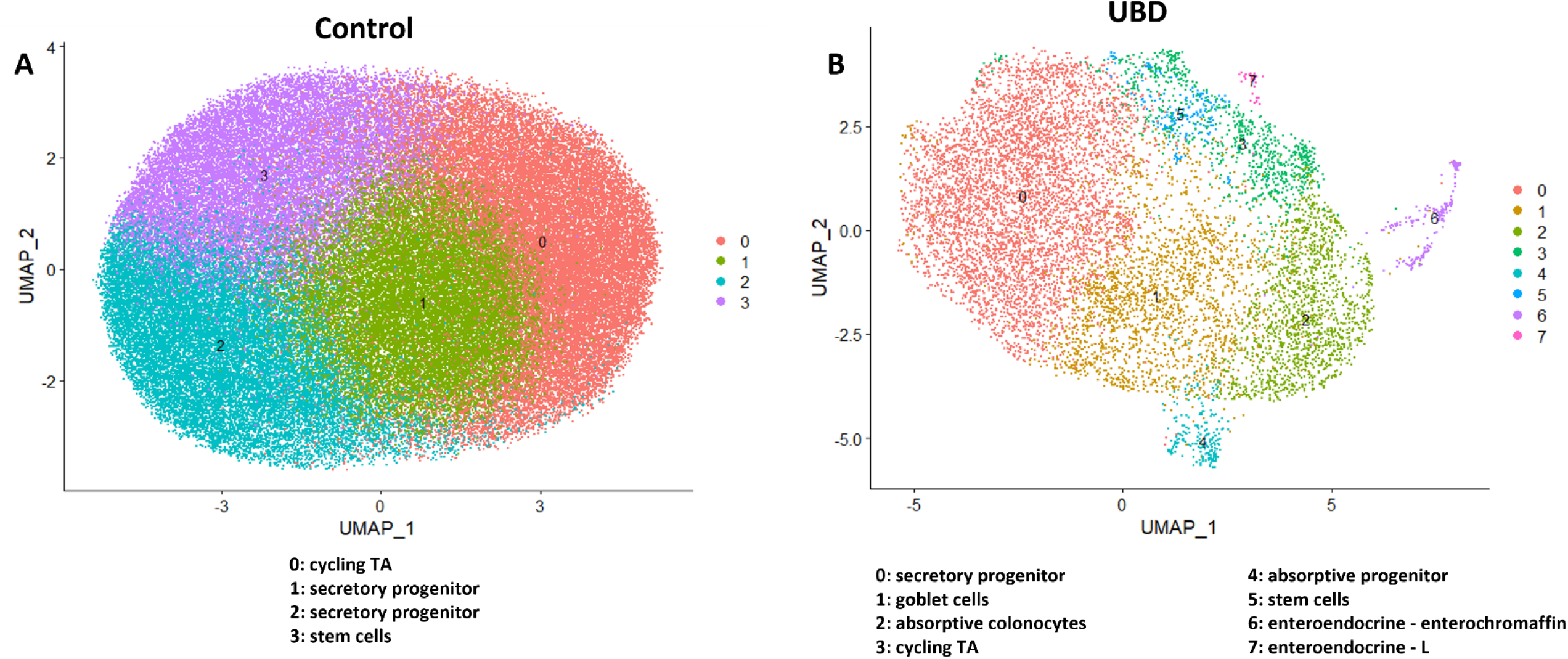
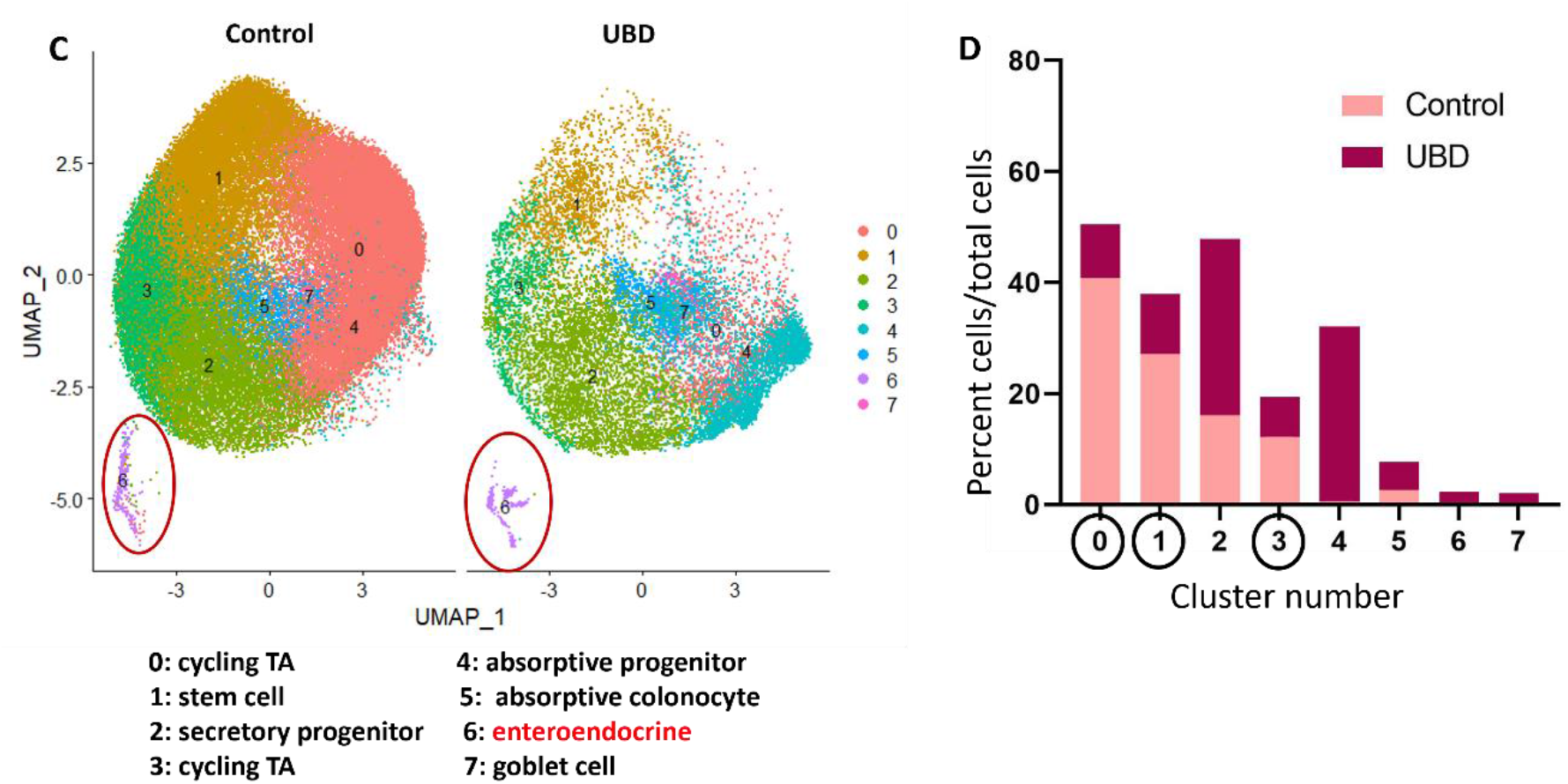
UMAPs of representative control and UBD treated colonoids show differentiated cells expand upon UBD treatment. **A)** Control and **B)** UBD treated colonoids were initially analyzed separately to determine differences between the biological controls. The representative control clusters show limited differential gene expression and cluster identities, primarily representing those residing in the deep crypt, while UBD treatment induces differentiation. **C)** Representative UMAP of merged control and UBD colonoids to perform direct comparison analysis between each cell cluster. **D)** Quantification of cells in each cluster shows that proliferative cell numbers decrease while differentiated cell numbers expand.

In contrast, UBD treated colonoids maintained in identical media, had a higher degree of cell type variability, as shown in the representative UBD treated colonoid UMAP. They grouped into clusters (**Dataset 1**) that included differentiated cell types and were identified as: secretory progenitors, goblet cells, absorptive colonocytes, cycling TA cells, absorptive progenitors, stem cells, enteroendocrine (EEC) – enterochromaffin cells and EEC – L cells **(Fig. 2B, right)**. This indicates that acute UBD exposure diversified the differentiated cell types in human colonoids. Since the three colonoid lines were from biologically unique donors, differential gene and clustering analysis was performed on each control and UBD treated colonoid set to first determine if there were major differences due to genetic variability. Similar clustering patterns and differentially expressed genes were seen in all three colonoid lines for control (mainly deep crypt cells) and UBD treated (mainly differentiated cells) conditions. Thus, control and UBD treated colonoid datasets were merged to directly compare cell proportions in each cluster (**Fig. 2C, Dataset 2**). There was an increase in all types of differentiated cells, but a decline in proliferative (stem and cycling TA) cells in the UBD treated colonoids (**Fig. 2D**).

To visualize differences within clusters between control and UBD treated colonoids, the datasets were merged, clustered, and split according to treatment. UBD treated cells (blue) do not entirely overlap with control cells (red), indicating UBD induced changes in gene expression patterns (**Supplemental Fig. 2**). Volcano plots of control and UBD treated colonoids for all 3 colonoid lines show significantly up (red) and down (blue) regulated genes by treatment (**Supplemental Fig 3**). In UBD treated colonoids, genes associated with secretory lineage (such as *TFF3, CHGA, PCSK1N, MUC2, FCGBP)* were upregulated, while cell cycle and mitochondrial genes were downregulated. In contrast, mitochondrial genes were upregulated in control colonoids. Taken together, these results further indicate that acute UBD treatment induces differentiation in human colonoids which leads to reciprocal shrinking of proliferative cells.

### UBD exposure induces EEC expansion in human colonoids

Disruption of EECs and hormones have been reported in various gut-related and systemic metabolic diseases (French et al., 1993; Jorsal et al., 2018; Keller et al., 2009). Among the cell type changes observed upon UBD treatment, there was an expansion in EECs (**Fig. 2C, cluster 6)**. Since EECs are rare and control colonoids did not show major biological variation when clustered independently, we merged the control and UBD treated colonoids for downstream analysis. We identified EECs by the expression of chromogranin A (*CHGA*) and chromogranin B (*CHGB*), both well characterized pan-EEC markers. Using feature plots to identify *CHGA* and *CHGB* expressing cells (**Fig. 3A**) and dot plots to quantify *CHGA* and *CHGB* expression (**Fig. 3B**), we found a 3.6-fold increase in total number of EECs, 4.4-fold increase of *CHGA*+ cells and 6.3-fold increase in *CHGB*+ cells (**Fig. 3C**) in UBD treated colonoids. We confirmed these increases by quantifying numbers of CHGA expressing cells via immunostaining (**Fig. 3D, E**). Additionally, detection of RNA oligos via RNAscope in-situ hybridization showed a significant increase in *CHGA*+ oligos per cell in UBD treated colonoids compared to controls (**Fig. 3F, G**). Therefore, EEC numbers significantly increase following acute UBD exposure in human colonoids.

**Fig 3.**
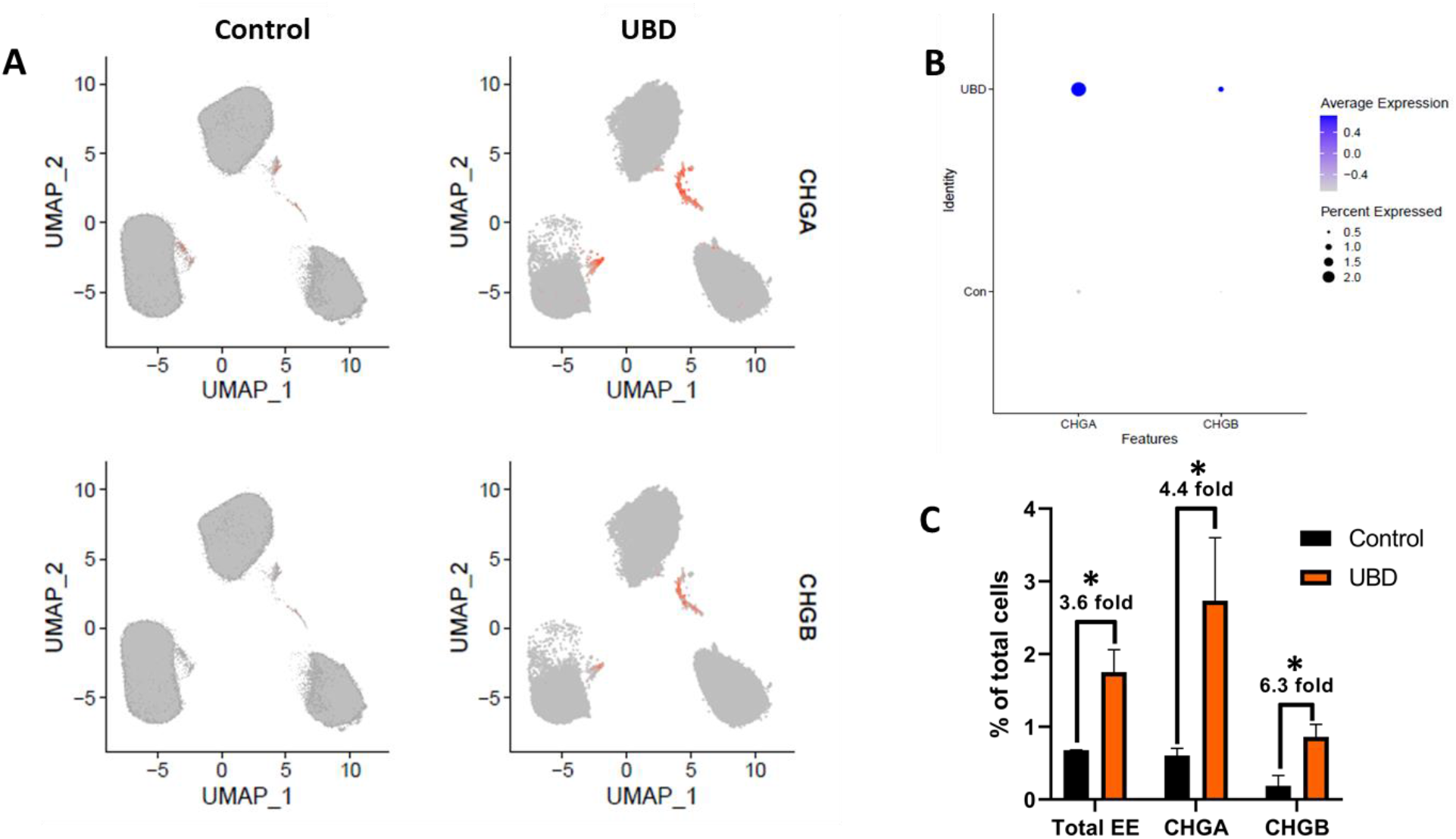
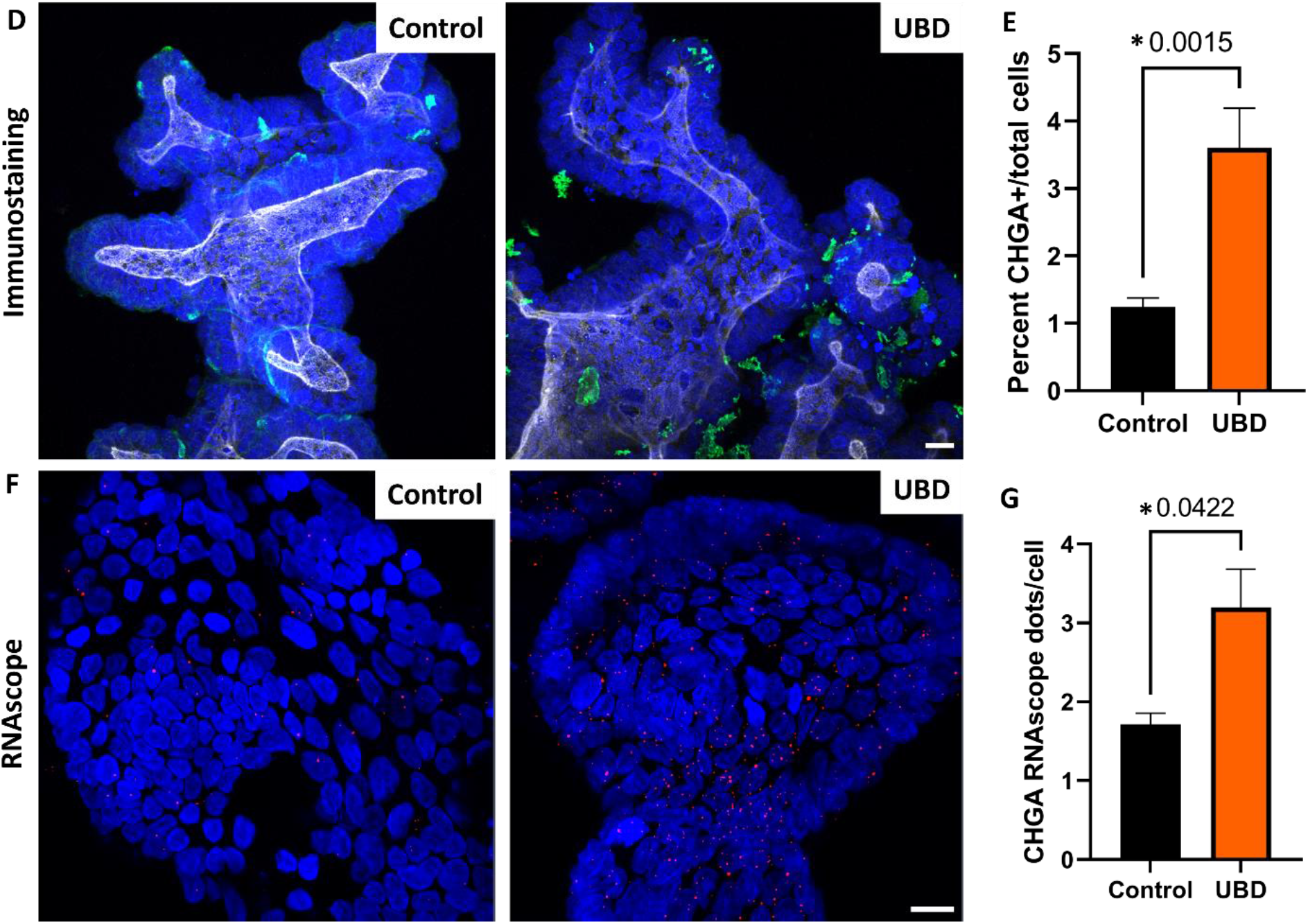
Enteroendocrine cells expand in colonoids exposed to UBD. **A)** Feature plot of pooled control (left) and UBD treated (right) colonoids (n=3) with cells differentially expressing CHGA (top) and CHGB (bottom) shown in orange. **B)** Dot plot of enteroendocrine markers CHGA and CHGB show increased expression in UBD treated colonoids. **C)** Quantification of total EECs and EEC markers CHGA and CHGB compared between the pooled control and UBD-exposed colonoids**. \***p-values=0.02 (total EE), 0.07 (CHGA), 0.03 (CHGB). **D)** Representative immunostaining of colonoids shows increase of CHGA expression upon UBD treatment. CHGA, green; phalloidin, white; nuclei, blue; scale bar = 20 µm. **E)** Quantification of percent CHGA+ cells in total cell populations in control and UBD treated colonoids. *p=0.0015; N≥5. **F)** Representative RNAscope fluorescence staining of CHGA in colonoids validates scRNA-seq. CHGA, red; nuclei, blue; scale bar = 20 µm. **G)** Quantification of RNAscope staining of CHGA dots per cell in total cell populations in control and UBD treated colonoids. *p=0.0422; N≥3. Data are presented as mean ± SEM.

### Serotonin producing-enterochromaffin cells and peptide YY-producing L cells expand upon UBD exposure

Enterochromaffin (EC) and L cells are the two most abundant EEC subtypes in the colon, however, there are at least 12 subtypes described in the small bowel and colon (Goldspink et al., 2018b). Both EC and L cells have previously been implicated in wound healing and regeneration in other organs, possibly due to their mitogenic potential (Lesurtel et al., 2006).

We further analyzed these two major EEC subtypes in our scRNA-seq datasets. EC cells were marked by expression of tryptophan hydroxylase 1 (*TPH1*), the enzyme that catalyzes the rate limiting step in synthesis of serotonin (5-HT) from tryptophan (Liu et al., 2021), while L cells were marked by expression of peptide tyrosine tyrosine (*PYY*) (Spreckley and Murphy, 2015). Feature plots of merged control and UBD treated colonoids showed a 5.4-fold expansion of EC cells and a 3.4-fold expansion of L cells following UBD treatment (**Fig. 4A-C**). A scatter plot of average gene expression in EECs by treatment shows that *PYY* is nearly absent in control colonoids but, like *CHGA*, is highly expressed in UBD treated colonoids (**Supplemental Fig. 4**).

**Fig 4.**
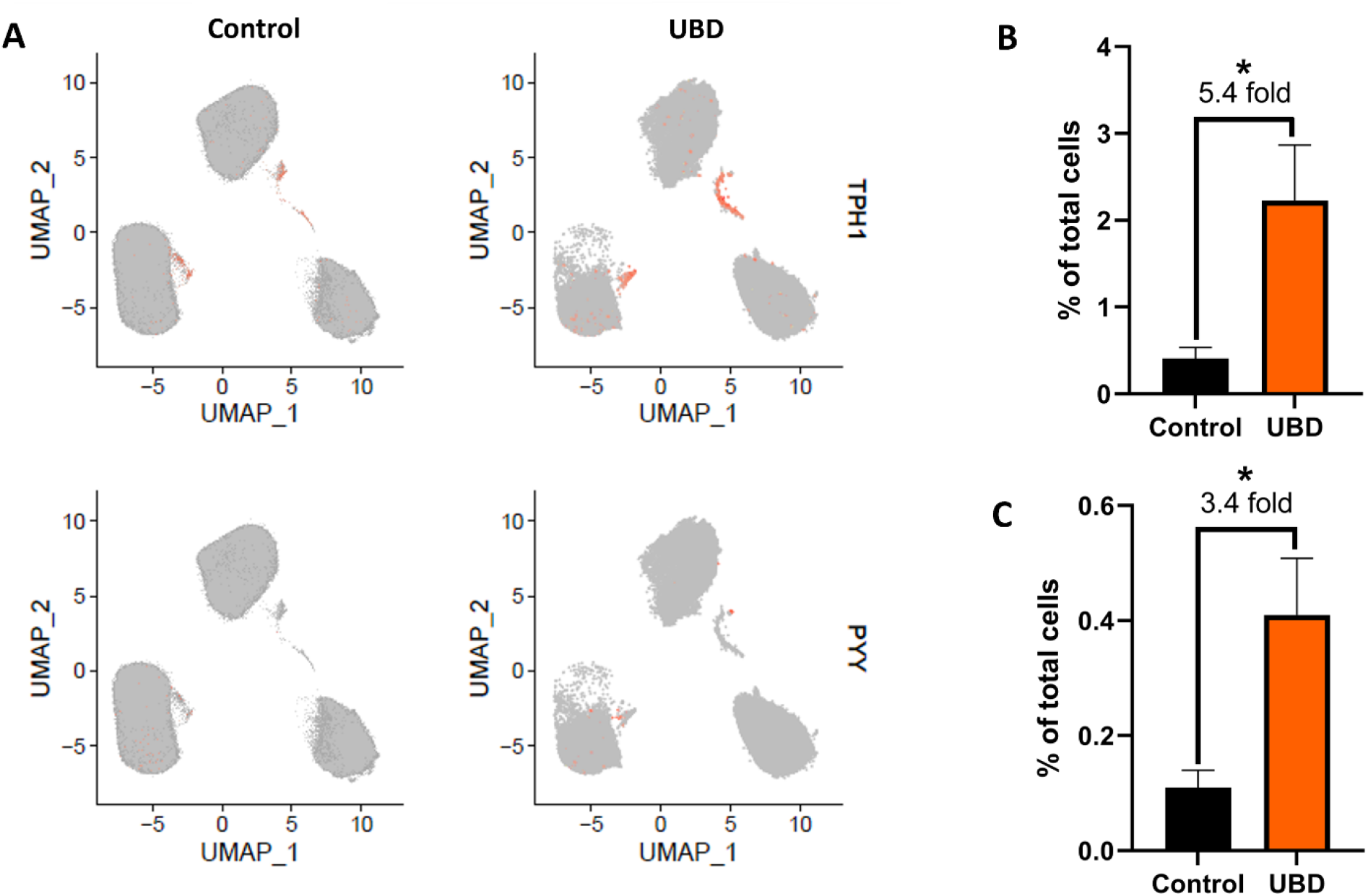
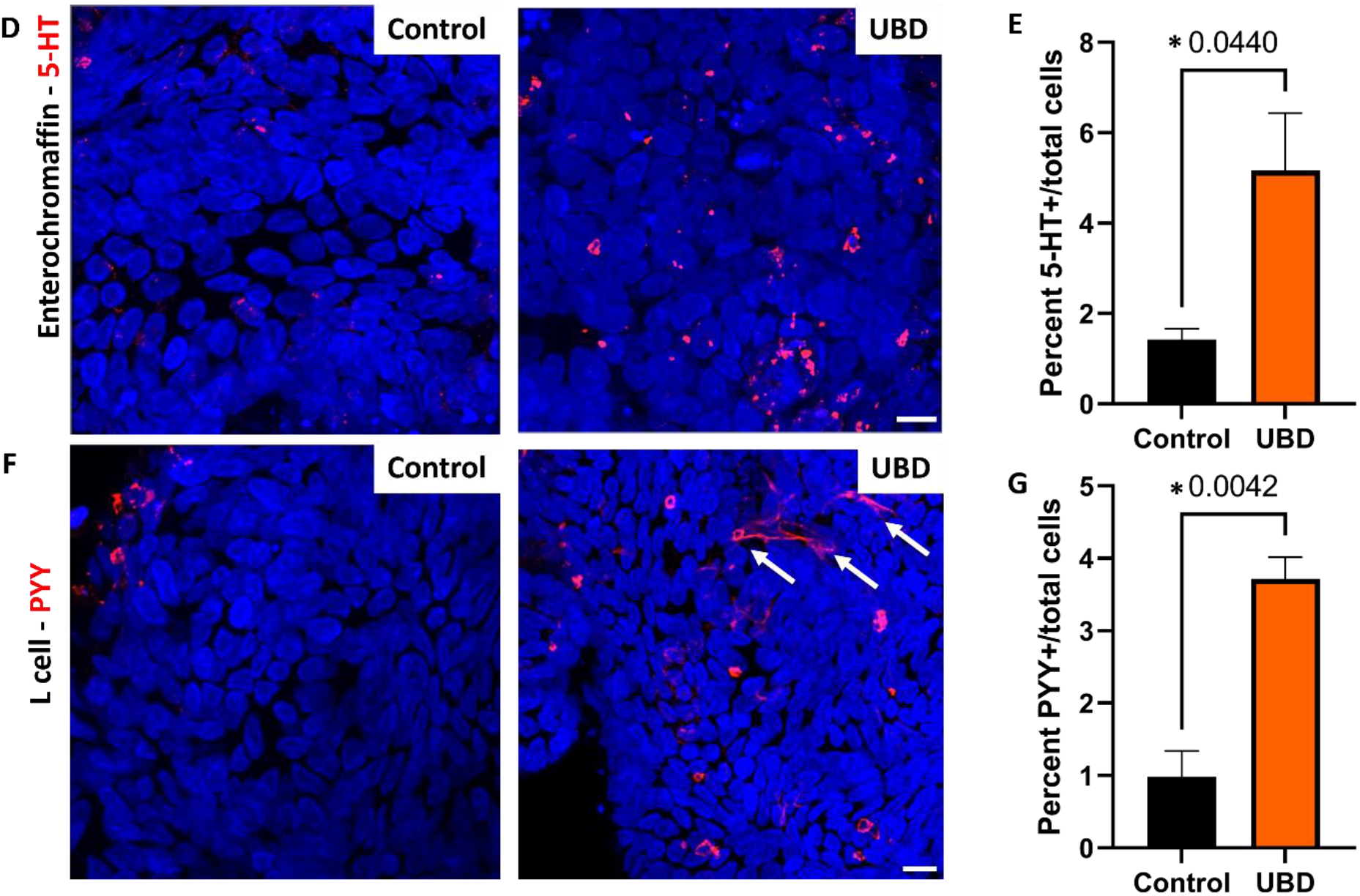
Enterochromaffin cells and L cells expand upon UBD treatment. **A)** Feature plot of pooled control (left) and UBD treated (right) colonoids (n=3) with cells differentially expressing TPH1 (top) and PYY (bottom) shown in orange. **B)** Quantification of cells differentially expressing TPH1 compared between the pooled control and UBD treated colonoids**. \***p=0.0497. **C)** Quantification of cells differentially expressing PYY compared between the pooled control and UBD treated colonoids**. \***p=0.0435. **D)** Representative immunostaining of colonoids shows increase of 5-HT upon UBD treatment. 5-HT, red; nuclei, blue; scale bar = 20 µm. **E)** Quantification of percent 5-HT+ cells in total cell populations in control and UBD treated colonoids. *p=0.044; N≥3. **F)** Representative immunostaining of colonoids shows increase of PYY in colonoids upon UBD treatment. White arrows indicate PYY+ basolateral extensions. PYY, red; nuclei, blue; scale bar = 20 µm. **G)** Quantification of percent PYY+ cells in total cell populations in control and UBD treated colonoids. *p=0.0042; N≥3. Data are presented as mean ± SEM.

We validated EC and L cell expansion upon acute UBD exposure by immunostaining for 5-HT (**Fig. 4D**) and PYY (**Fig. 4F**), respectively. Quantification of 5-HT showed a significant increase in EC cells in UBD treated colonoids compared to controls (**Fig. 4E**). Similarly, the proportion of PYY expressing L cells were found to significantly increase in UBD treated colonoids when compared to controls (**Fig. 4G**). Interestingly, we observed PYY+ basal extensions (**Fig. 4F**, white arrows) from UBD treated L cells that were absent in control colonoid L cells.

### Sub-clustering EECs at higher resolution reveals new subtypes in UBD treated colonoids

To further determine the spectrum of EECs that results from acute UBD exposure, we re-analyzed the EEC clusters from the combined control and UBD treated colonoids. The EEC clusters were sub-clustered at a higher resolution of 1 instead of 0.4, then identified by comparing the differentially expressed genes (**Dataset 3**) with published EEC datasets (Beumer et al., 2020; Gehart et al., 2019). This resulted in the control colonoid EECs separating into three clusters identified as: EEC progenitors, X cells and EC cells (**Fig. 5A, left UMAP**). The UBD treated EECs sub-clustered into eight distinct clusters identified as: EEC progenitors, X cells, EC cells, N cells, K cells, L cells and enterochromaffin-late cells (**Fig. 5A, right UMAP**). Sub-clustering also identified two subtypes of L cells in the UBD treated colonoids. A small group of L cells expressed both GCG (proglucagon) and PYY (circled in purple) while a larger group of L cells expressed only PYY (circled in red) (**Fig. 5B**). These results indicate that UBD treatment induces differentiation of EEC subtypes that are not found in control colonoids.

**Fig 5.**
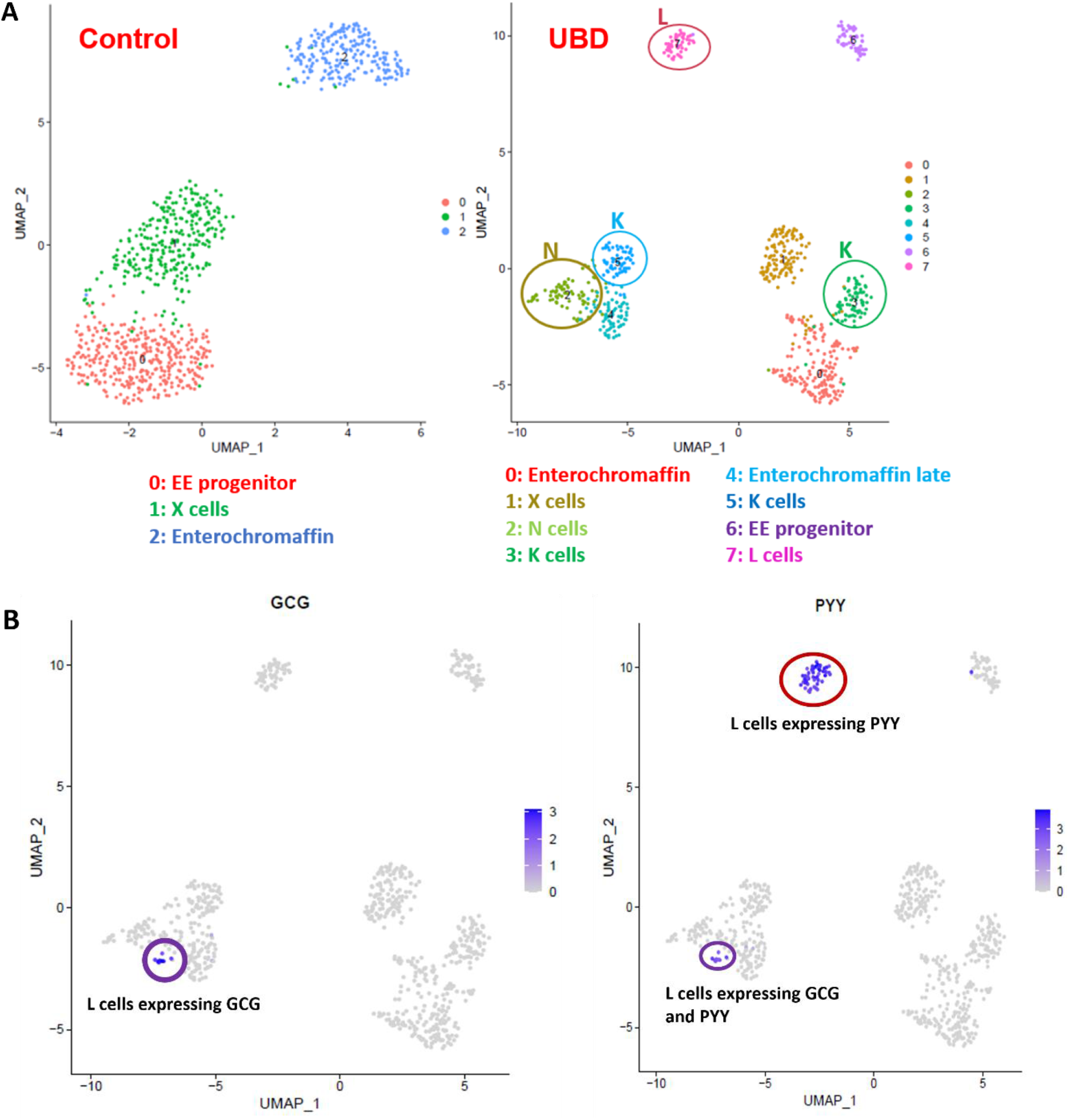
Sub-clustering at higher resolution reveals UBD induces increased enteroendocrine cell subtype differentiation in colonoids. **A)** Representative UMAP of EECs in control and UBD treated colonoids sub-clustered at a higher resolution and identified. UBD treatment induces further differentiation into EEC subtypes not seen in control colonoids. **B)** Feature plots of UBD treated EECs separated by GCG or PYY differential expression. Some L cells express both GCG and PYY (purple circle), but a subset of L cells express only PYY (red circle).

### UBD exposure changes the gene expression profile of EECs

We evaluated the transcriptome of the newly formed UBD-induced EECs compared to control colonoid EECs. We compared gene profiles of the three common EEC clusters (progenitors (blue), X cells (green), and EC cells (red)) between control and UBD-treated colonoids (**Fig. 6A**). The most notable differences were seen in the EEC progenitor clusters. The topmost expressed genes in the EEC progenitor cluster in control (**Fig. 6A**, cluster 0, blue box) are associated with secretory cells. In contrast, genes in EEC progenitors in UBD treated cells (**Fig. 6A**, cluster 6, blue box) are associated with the cell cycle.

**Fig 6.**
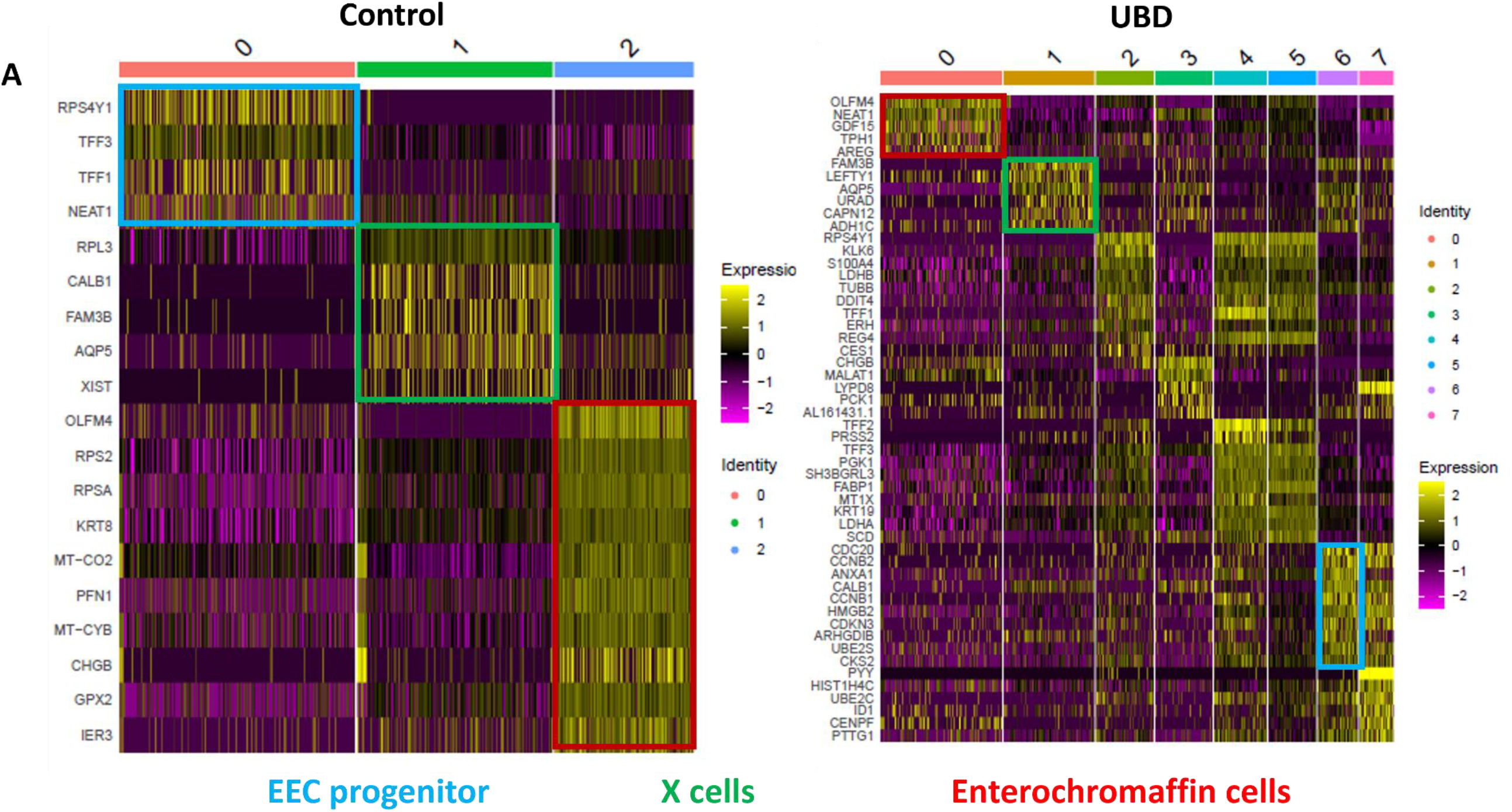
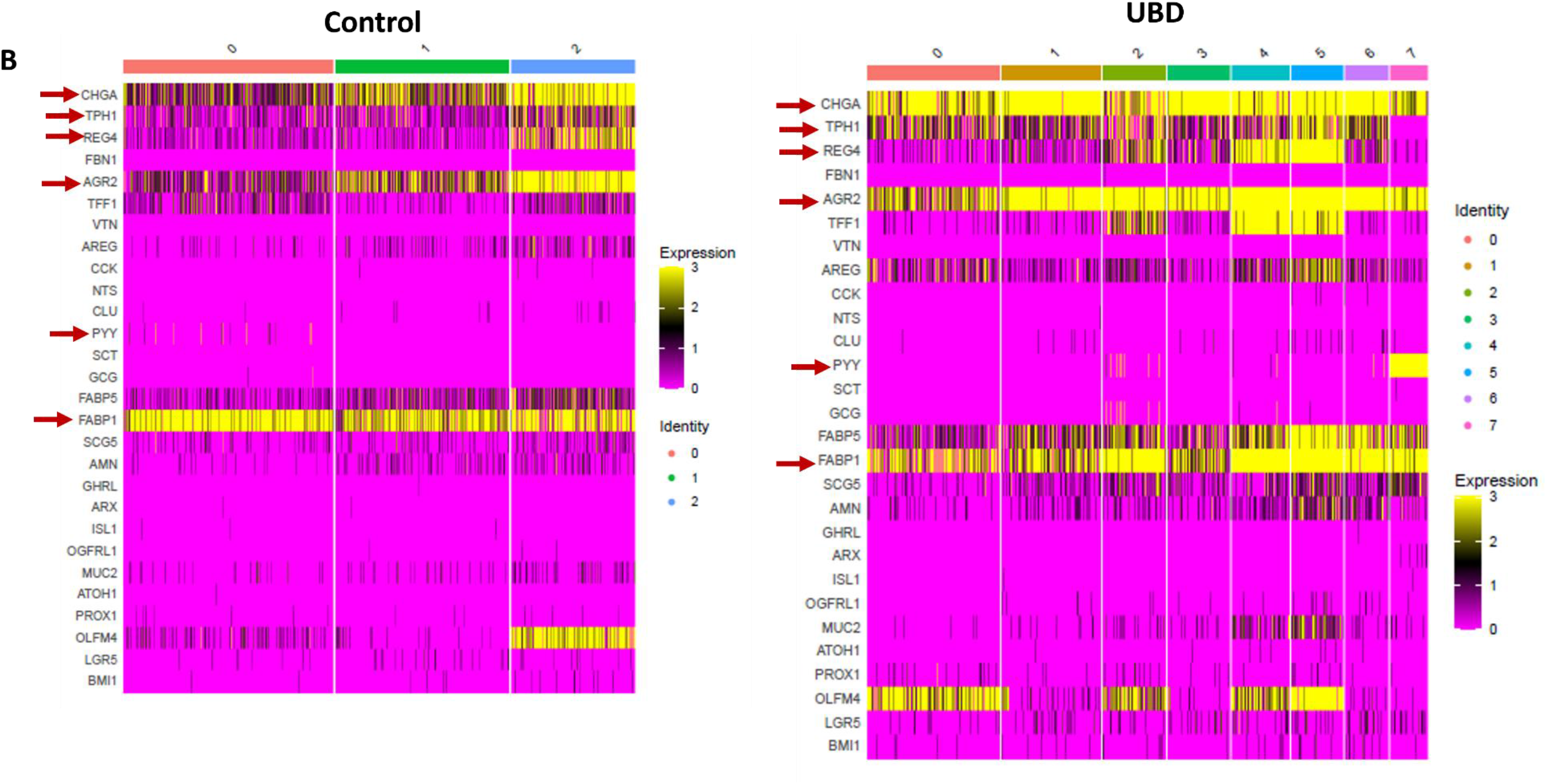
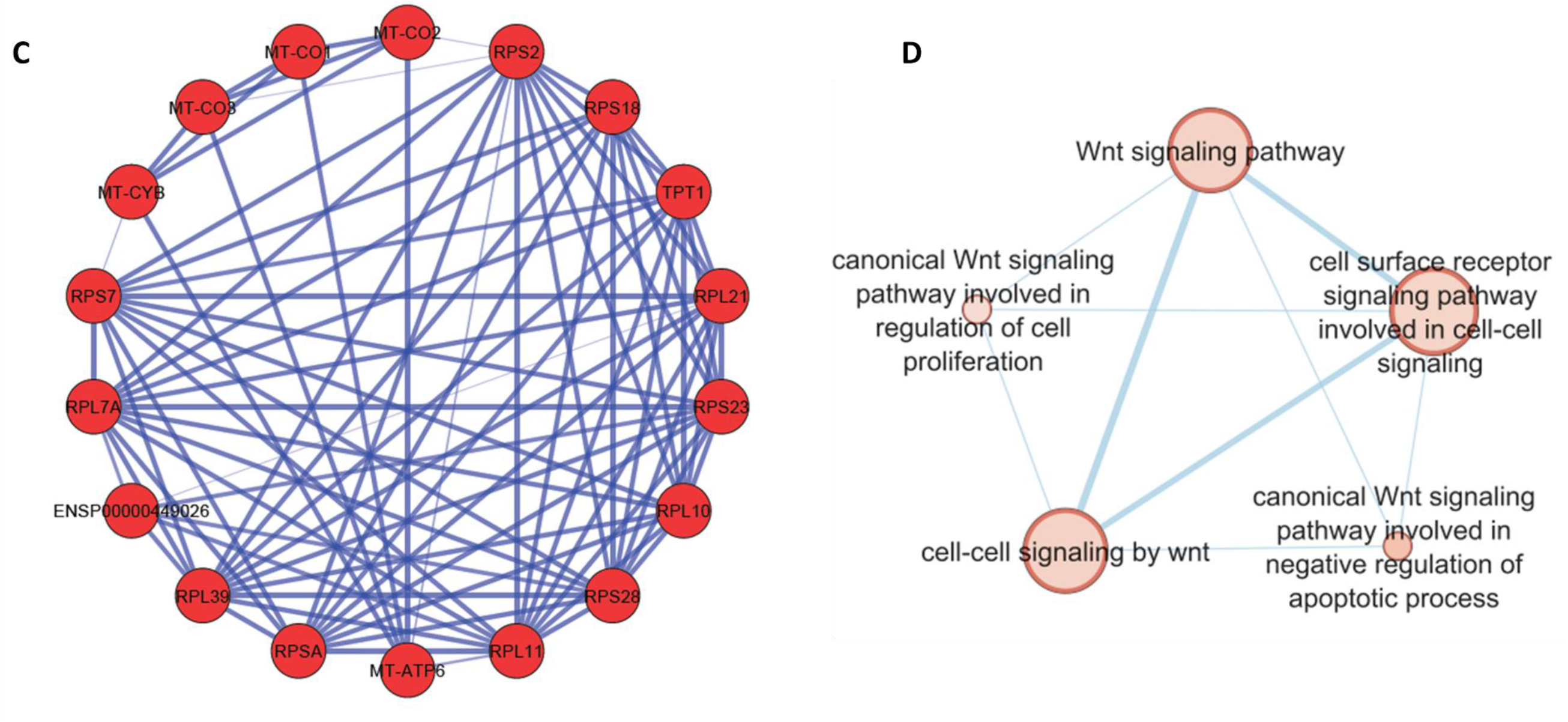
UBD exposure changes the gene expression profile of enteroendocrine cells. **A)** Heatmap of pooled EECs from control colonoids (left) and UBD treated colonoids (right) comparing the gene profiles of EEC progenitors, X cells, and enterochromaffin cells between the two conditions. Top upregulated genes in EEC progenitors in control (cluster 0, blue box) are associated with secretory cells while the EEC progenitors in UBD treated (cluster 6, blue box) are associated with cell cycle. **B)** Heatmap of selected EEC genes show increased expression of CHGA, TPH1, REG4, AGR2, PYY, and FABP1 in the UBD treated (right) compared to control colonoids (left). **C)** Network analysis of top upregulated genes in UBD treated EECs shows increased interactions between mitochondrial and ribosomal genes. **D)** Enrichment map of UBD treated EECs shows upregulation of multiple pathways related to WNT signaling.

Comparing relative expression amounts of select EEC-associated genes, we found that *CHGA, TPH1, REG4, AGR2, PYY*, and *FABP1* had increased expression in the UBD treated colonoids compared to control colonoids (**Fig. 6B**). We performed differential gene analysis on EECs from UBD-treated colonoids to further characterize the transcriptomic changes. Over 700 genes were differentially expressed in EECs upon UBD treatment, of which 67% were upregulated. Network analysis of top upregulated genes in UBD treated EECs showed interactions between mitochondrial and ribosomal genes (**Fig. 6C**). This indicates an increase in metabolism and protein synthesis required for cell proliferation and differentiation. We performed pathway analysis in gProfiler using the genes that were upregulated in UBD treated EECs. Upregulated pathways from gProfiler were imported into Cytoscape for enrichment analysis. The enrichment map of UBD treated EECs shows upregulation of multiple pathways related to WNT signaling (**Fig. 6D**).

### EEC differentiation occurs via a non-canonical mechanism in UBD treated colonoids

Based on our analysis indicating that EEC hyperplasia in UBD treated colonoids may be related to upregulated WNT signaling, we sought to understand the mechanism of EEC differentiation in the context of UBD induced injury. *NEUROG3* and *NEUROD1* are the early and late transcription factors that drive EEC differentiation in mammals (Li et al., 2011). Both transcription factors are considered WNT-independent, which is contrary to our network analysis of EECS in UBD treated colonoids. Neither *NEUROG3* nor *NEUROD1* expression was greatly increased upon UBD treatment compared to control, as depicted in the dot plot analysis of the scRNA-seq datasets (**Fig. 7A**).

**Fig 7.**
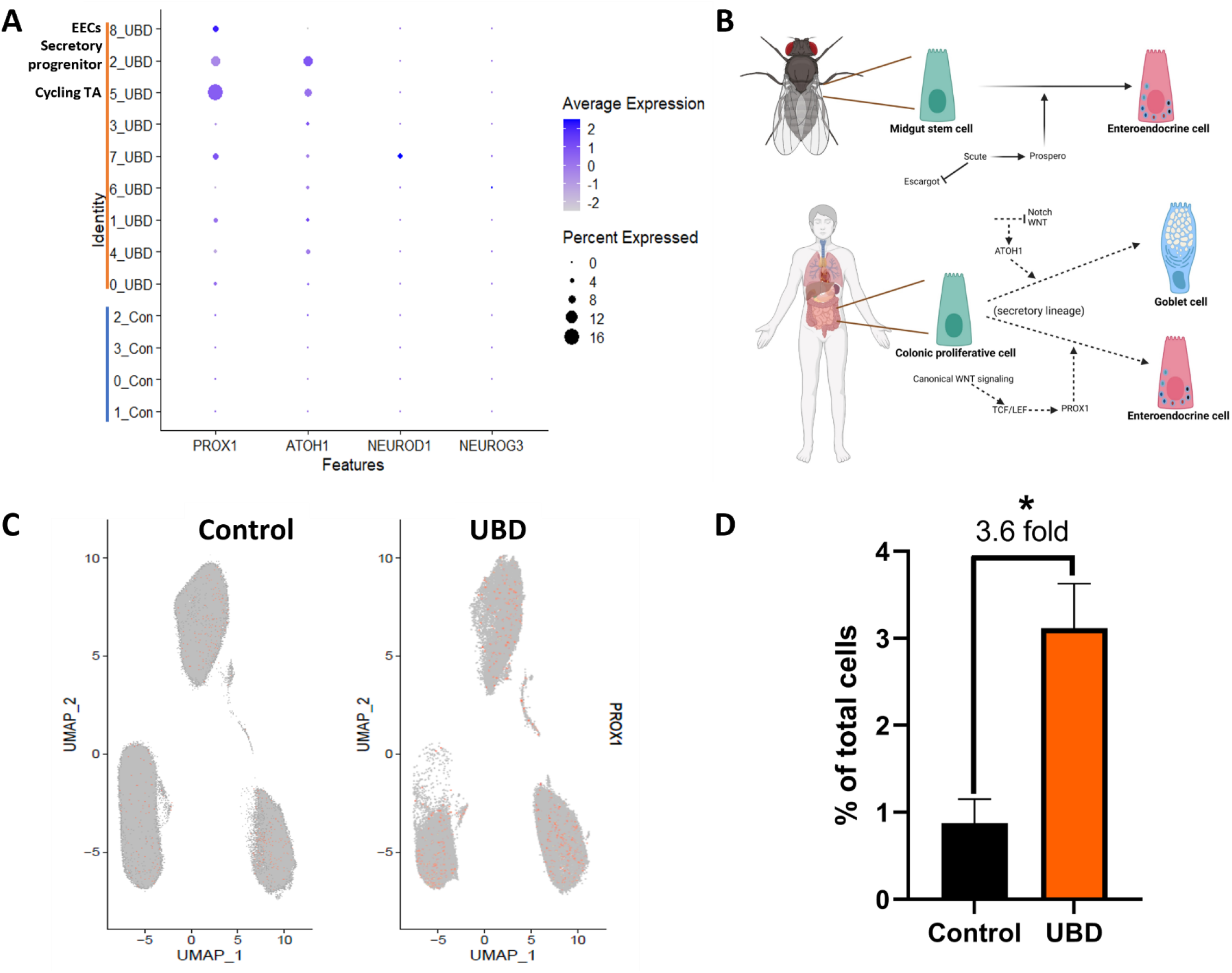
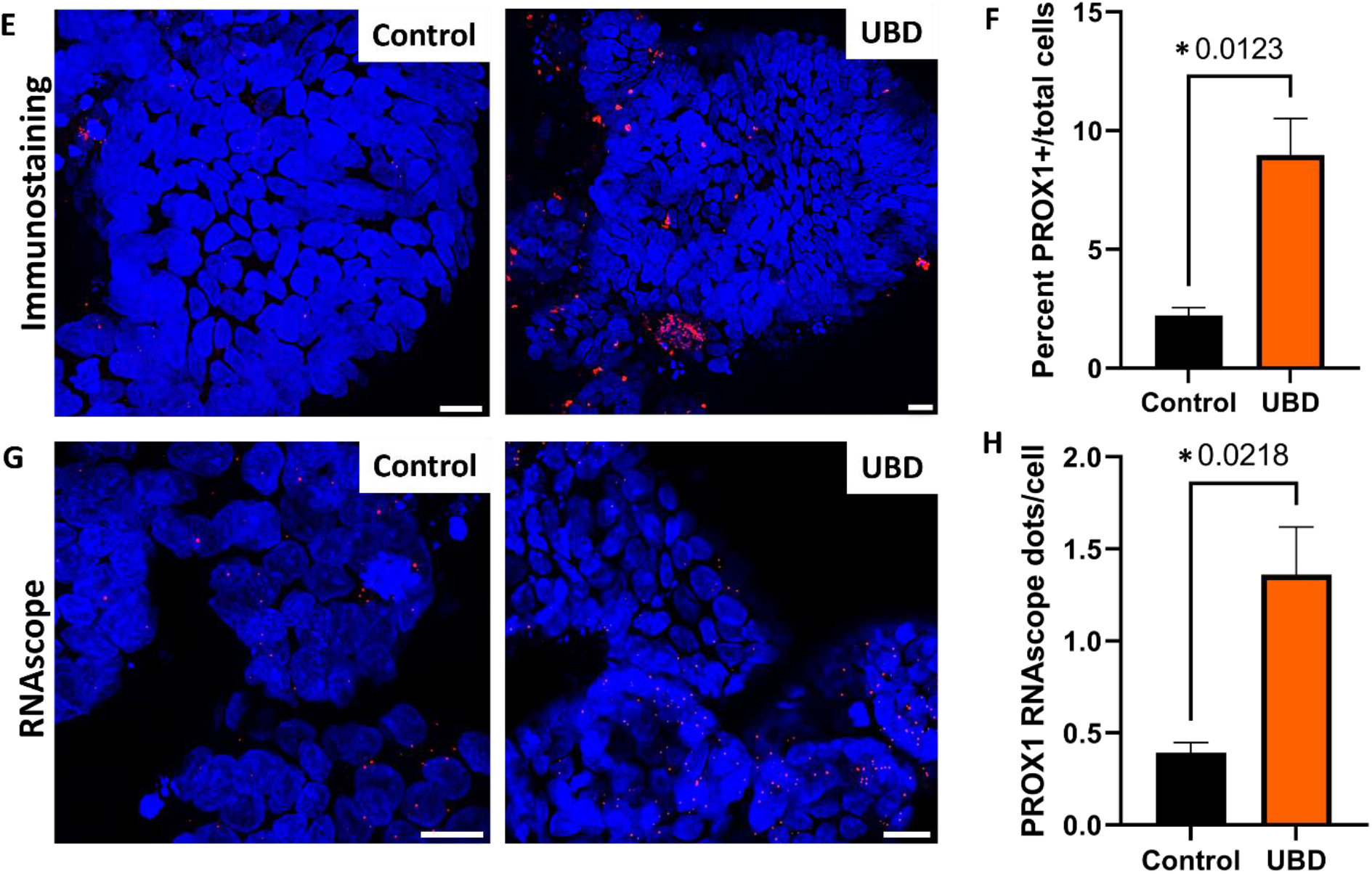
PROX1 is associated with increased EEC differentiation in UBD treated colonoids. **A)** Dot plot of transcription factors PROX1, ATOH1 (goblet cells), NEUROD1 (EECs), and NEUROG3 (EECs) shows that only PROX1 is upregulated upon UBD exposure in cycling TA, secretory progenitor, and EECs. **B)** Schematic showing EEC differentiation is regulated by Prospero in *Drosophila* (top) and possibly PROX1, the human homolog of Prospero, in human colon (bottom). Created in *Biorender.* **C)** Feature plot of cells differentially expressing PROX1 (orange) in pooled control and UBD treated colonoids (n=3). **D)** Quantification of cells differentially expressing PROX1 compared between the pooled control and UBD treated colonoids**. \***p=0.0179 **E)** Representative immunostaining of colonoids shows increase of PROX1 upon UBD exposure. PROX1, red; nuclei, blue; scale bar = 20 µm. **F)** Quantification of percent PROX1+ cells in total cell populations in control and UBD treated colonoids. *p=0.0123; N≥3. **G)** Representative RNAscope fluorescence staining of PROX1 in colonoids validates scRNA-seq. PROX1, red; nuclei, blue; scale bar = 20 µm. **H)** Quantification of RNAscope staining of PROX1 dots per cell in total cell populations in control and UBD treated colonoids. *p=0.0218; N≥3. Data are presented as mean ± SEM.

Thus, we searched for a possible WNT-dependent transcription factor that may be driving UBD-induced EEC differentiation. In *Drosophila*, the transcription factor Prospero drives differentiation of midgut stem cells into EECs (Zeng and Hou, 2015). The mammalian homolog of Prospero is PROX1 and is WNT-dependent (Cha et al., 2018). Therefore, we hypothesized that UBD treatment leads to activation of WNT signaling and upregulates PROX1 expression, driving EEC differentiation from secretory progenitor cells (**Fig. 7B**). From our scRNA-seq analysis, there was a significant increase in *PROX1*-expressing cells in UBD treated colonoids compared to control colonoids **(Fig. 7C, D)**. Additionally, in UBD treated colonoids, *PROX1* expression was significantly increased in cycling TA and secretory progenitor cells (**Fig. 7A**). This PROX1 increase was validated via immunostaining (**Fig. 7E, F**) and RNAscope (**Fig. 7G, H**). These results suggest EEC differentiation upon UBD treatment might be independent of the canonical transcription factors *NEUROG3* and *NEUROD1.* Instead, it is dependent on the WNT-dependent transcription factor *PROX1*. Overall, these results show that environmental heavy metal-induced injury in the gut can be modeled by human colonoids, and that EEC differentiation is an acute response to this environmental injury.

## DISCUSSION

Gut exposure to environmental toxins and heavy metals has been linked to intestinal and systemic diseases, as well as inhibition of intestinal epithelial healing (Chiu et al., 2020; Wagner et al., 2011; Wales and Davies, 2015). However, transcriptomic changes within intestinal epithelial cells due to these environmental toxicants are largely unknown. UBD was used to model environmental exposure of ^238^U to human colonoids (a simplified, yet physiologically relevant intestinal model). Droplet based scRNA-seq demonstrated that acute UBD injury has a distinct effect on the gut. EECs expanded and increased differentiation into *de novo* EEC sub-types. We also observed enrichment of WNT signaling, but a decrease in proliferative cells, and identified a potential non-canonical mechanism of EEC differentiation upon UBD-induced injury.

The mammalian intestinal epithelium regenerates continuously by a proliferative pool of stem and transient amplifying cells (Rees et al., 2020). These proliferative cells give rise to various mature cell types required for an intact and properly functioning epithelial barrier. Thus, a decrease in proliferative cells, as seen with acute UBD exposure, hampers the ability of the gut to repair its epithelium with potentially severe downstream consequences. Impairment of epithelium regeneration augments access of luminal toxins, antigens, and microbes to deeper layers of the mucosa. This can lead to recruitment of immune cells and subsequent inflammation (Zundler et al., 2019).

Another major observation was the expansion of hormone-producing EECs, which are typically rare and make up ∼1% of the total gut epithelium (Sternini et al., 2008). The intestine is the largest endocrine organ in humans (Buchan, 1999) and communicates with the brain via EECs and the enteric nervous system (Kuwahara et al., 2020; Latorre et al., 2016). Gut-brain communication is partially attributed to EEC neuropods that directly synapse with enteric nerves (Bohorquez et al., 2011; Latorre et al., 2016). Therefore, modulation of the intestine via environmental toxicants can indirectly affect systemic health.

In this study, UBD induced a 5-fold expansion of 5-HT-producing EC cells. In the gut, 5-HT stimulates motility and secretion (Gershon and Tack, 2007; Pithadia and Jain, 2009), enhances mucosal immunity and has been implicated in wound healing. Peripheral 5-HT can circulate systemically via platelets and act as a mitogen in injured organs. In murine models of liver injury, 5-HT was able to promote liver regeneration after partial hepatectomy (Lesurtel et al., 2006), while mice lacking TPH1 were unable to fully regenerate their liver. These findings suggest that EECs can undergo adaptive reprogramming during injury and provide insights for potential new regenerative therapeutics in the gut. Despite the beneficial effects of 5-HT, high levels can be detrimental. As a vasoconstrictor, sustained 5-HT release elevates blood pressure with risk of severe fluctuations in hypertensive individuals (Amstein et al., 1988), decreases bone mineral density (Bliziotes, 2010) and dysregulates metabolic processes like gluconeogenesis and lipolysis which may predispose individuals to diabetes and obesity (Yabut et al., 2019).

Acute UBD injury also led to a 3-fold expansion in L cells, the second most abundant EEC subtype in the colon. The major peptides secreted by L cells: PYY, glucagon-like peptide 1 (GLP-1) and glucagon-like peptide 2 (GLP-2), are associated with intestinal motility, protection against inflammation and regeneration (Drucker, 2018; Drucker et al., 1996; Lebrun et al., 2017). Based on the known function of L cell peptides, this suggests that UBD-induced L cell expansion may be beneficial by stimulating mitogenesis essential for regeneration.

We identified EEC subtypes (N and K cells) in the UBD treated EEC sub-clusters that were not present in control colonoids. In homeostatic conditions, K cells are primarily found in the small bowel (Rindi et al., 2004). Their appearance in the colon is unexpected. K cells produce glucose-dependent insulinotropic peptide (GIP), which inhibits gastric acid secretion and gastrointestinal motility (Cho and Kieffer, 2010). It is counter-intuitive to slow gut motility, as we would expect an increase in gut motility and secretion to flush away toxicants from the gut. Again, this suggests that we do not have a full understanding of EECs and the functional roles of their specific hormones and peptides. Neurotensin produced by N cells stimulates colonic regeneration, gut motility (Evers et al., 1992; Rock et al., 2022) and can facilitate exit of toxicants from the body. This suggests that *de novo* differentiation of N cells during UBD exposure may be beneficial.

From our results and previous studies, we see that EECs can be dynamically altered during injury. There is evidence that implicates EECs as facultative stem cells that promote plasticity and regeneration (Johnson et al., 2022; Rindi et al., 2004). However, EEC expansion can also contribute to the rise of small intestinal neuroendocrine tumors (Sei et al., 2020). EECs can revert to stem cells following injury with high intensity radiation (12 Gy) that causes massive cell death and ablation of crypt-based stem cells (Yan et al., 2017). On the contrary, UBD induced injury is mild and does not cause massive cell death. Instead, UBD injury induces changes to lineage differentiation, particularly expansion of *de novo* and mature EECs. These opposing observations suggest EEC response to injury might depend on severity of damage, further exemplifying the plasticity of these cells.

Downstream analysis of our scRNA-seq datasets found enrichment of WNT signaling pathways in UBD-induced EECs. However, neither *NEUROG3* (Gehart et al., 2019; Li et al., 2011), nor *NEUROD1* (Li et al., 2011; Schonhoff et al., 2004), the canonical transcription factors that drive EEC differentiation (Li et al., 2011; Schonhoff et al., 2004), are WNT dependent. Instead, our scRNA-seq data found *PROX1,* a WNT dependent transcription factor (Karalay et al., 2011), was greatly upregulated in cycling TA and secretory progenitor cells in UBD treated colonoids. *PROX1* is the mammalian homolog of Prospero, the *Drosophila* transcription factor (Karalay et al., 2011; Zeng and Hou, 2015) that drives differentiation of midgut stem cells into EECs (Zeng and Hou, 2015). Since PROX1 is a WNT dependent transcription factor and given that WNT signaling was enriched in UBD-induced EECs, we hypothesize that injury from toxicants leads to increased WNT signaling, which upregulates PROX1 and drives secretory progenitor cells into non-canonical EEC differentiation.

The UBD used in this study was obtained from the attic of a structure located on Laguna Pueblo, less than 1 km from the edge of a now dormant, 3000 acre open-pit uranium mine. A caveat to this study is the lack of knowledge on the causal factors in the UBD, such as physical and chemical interactions of uranium or other heavy metals and particulates. Importantly, samples did not generate measurable radiation and the uranium in the particulate dust was primarily non-fissile. Since this study focused on assessing transcriptional and cell differentiation effects between control and UBD treated colonoids, extensive investigation into the physicochemical properties of UBD would not enable further insights into this putative toxicant.

In summary, using a real-world environmental toxicant, we showed that human colonoids develop a pathophysiological response that shifts epithelial cell lineages. Proliferative cells are decreased while secretory lineage differentiation is increased, most notably EECs. Importantly, EEC plasticity leads to the development of EEC subtypes that are not present in control colonoids. Further investigation is required to determine if this expansion is mostly beneficial or detrimental, since enterohormones can provide both regenerative and systemic dysfunction effects. Understanding changes to EEC differentiation in injury and regeneration will provide the basis for better understanding of systemic changes and dysfunction that can originate from enterohormones in the gut.

## METHODS

### Tissue collection and colonoid generation

Human colonoid studies were reviewed and approved by the Johns Hopkins University School of Medicine Institutional Review Board (IRB# NA_00038329) and University of New Mexico Institutional Review Board (IRB# 18-171). Colonic biopsies were obtained from healthy individuals undergoing screening colonoscopies who had given informed written consent. Colonic crypt isolation and colonoid generation were prepared as previously reported (In et al., 2016; Jung et al., 2011). Briefly, biopsy tissue was minced, washed several times in freshly prepared cold chelating solution (CCS; 5.6mM Na2HPO4, 8mM KH2PO4, 96.2mM NaCl, 1.6mM KCl, 43.4mM sucrose, 54.9mM D-sorbitol, and 0.5mM DL-dithiothreitol) and incubated 1 hour at 4°C in CCS with 10 mM EDTA on an orbital shaker. Isolated crypts were resuspended in Matrigel (Corning, Tewksbury, MA) and 30 ul droplets were plated in a 24-well plate (Corning). After polymerization at 37°C, 500 ul of organoid expansion media was added for 2 days (Advanced Dulbecco’s modified Eagle medium/Ham’s F-12 (ThermoFisher, Waltham, MA), 100 U/mL penicillin/streptomycin (Quality Biological, Gaithersburg, MD), 10 mM HEPES (ThermoFisher), and 1X GlutaMAX (ThermoFisher), with 50% v/v WNT3A conditioned medium (ATCC CRL-2647), 15% v/v R-spondin1 conditioned medium (cell line kindly provided by Calvin Kuo, Stanford University), 10% v/v Noggin conditioned medium (cell line kindly provided by Gijs van den Brink, Tytgat Institute for Liver and Intestinal Research), 1X B27 supplement (ThermoFisher), 1mM N-acetylcysteine (MilliporeSigma, Burlington, MA), 50 ng/mL human epidermal growth factor (ThermoFisher), 10 nM [Leu-15] gastrin (AnaSpec, Fremont, CA), 500 nM A83-01 (Tocris, Bristol, United Kingdom), 10 μM SB202190 (MilliporeSigma), 100 mg/mL primocin (InvivoGen, San Diego, CA), 10 μM CHIR99021 (Tocris), and 10 μM Y-27632 (Tocris)). After 2 days, the media (without CHIR99021 and Y-27632) was replaced every other day. Colonoids were passaged every 10-14 days by harvesting in Cultrex Organoid Harvesting Solution (Bio-Techne, Minneapolis, MN) at 4°C with shaking for 45 min. Colonoids were fragmented by trituration with a P200 pipet 30-50 times, collected and diluted in Advanced DMEM/F12 and centrifuged at 300 xg at 4°C for 10 min. The pellet was resuspended in Matrigel and plated as described for crypt isolation. All colonoid cultures were maintained at 37°C and 5% CO_2_. Unless noted, colonoid lines have been passaged >20 times.

### Brightfield imaging

Colonoids plated in Matrigel in 24 well plates were imaged during experiments on a Zeiss Axio Observer A1 inverted microscope (Zeiss, Oberkochen, Germany) with images captured on CellSense imaging software (Olympus, Tokyo, Japan). Images were viewed and processed using OlyVia (Olympus).

### Uranium-bearing dust analysis

UBD was obtained from the attic of a structure on indigenous lands that lie adjacent to the 3000-acre Jackpile mine in New Mexico. Samples of the collected dust were sieved using a 20 µM filter. Transmission electron microscopy (TEM) analyses of the dust were carried out on the <20 µM size fraction. Particulates from this fraction, suspended in acetone, were deposited onto holey carbon film supported on Cu TEM grids using pipettes. For the TEM analysis, five 50 x 50 µM TEM grid squares were studied in detail. The samples were air dried and then analyzed using a JEOL 2010F FEG TEM/STEM (JEOL USA, Peabody, MA) operating at 200 kV. The samples were imaged using a variety of different TEM and scanning electron transmission microscopy (STEM) techniques, including bright-field TEM and high-angle annular dark-field STEM imaging. Uranium-bearing particulates were identified on the JEOL 2010F by energy dispersive X-ray spectroscopy (EDS) using an Oxford Instruments AZtecEnergy EDS system (Oxford Instruments, United Kingdom) coupled to an Oxford X-Max 80 mm^2^ silicon drift detector (SDD). The detailed distribution of U-bearing phases and their association with other minerals was obtained using STEM EDS spectral X-ray mapping.

### Uranium-bearing dust sample mineralogy

Mineralogy of the <20 µM size fraction in the dust is based on an analysis of ∼1000 individual mineral grains or grain clusters and is shown in **Supplemental Figure 1**. This fraction consists of a variety of minerals that are commonly present in aeolian dust in arid environments including quartz, feldspar, micas/clay minerals, calcite, and gypsum. Rare minerals include baryte, rutile, and iron oxides such as hematite and magnetite. In addition to these minerals, extensive searching of the TEM grids revealed the presence of submicron grains of U-bearing minerals. In two of the five 50 x 50 µM TEM grid square areas, no U-bearing minerals were found, but in the three other grid squares, several grains were found, often in close spatial association. TEM and STEM images of two examples of these U-bearing grains are shown in **Supplemental Figure 1.** Of relevance, 1) no U-bearing grains were found with grain sizes greater than 0.5 µM, 2) the grains always occur in clusters intimately mixed with other minerals – no individual isolated grains were found, and 3) the individual U-bearing minerals can be as small as 50 nM in size or less. Estimating the abundance of these grains is very challenging from TEM data, but our estimate is ∼0.01%. The mass fraction is significantly lower, because the grains are all submicron, a characteristic that is unique to the U-bearing minerals.

### Treatment and preparation of single cells from 3D colonoids for single cell RNA-sequencing

Colonoids used for sequencing are described in **Supplemental Table 1.** Colonoids in Matrigel were treated overnight (18 h) with 50 µg/ml UBD. Next day, colonoids were removed from Matrigel by incubation with Organoid Harvesting Solution at 4°C for 45 min, washed in Advanced DMEM/F12 and pelleted by centrifugation at 300 xg for 10 min. Colonoids were resuspended in 500µl TrypLE enzyme solution (ThermoFisher), then transferred to a 24 well plate and spun at 600 rpm at 37°C for 45 min in a spinoculator (Mixer HC, USA scientific) to digest into single cells. Single cells were washed in 10 ml of Advanced DMEM/F12, pelleted again, resuspended in 1 ml of organoid expansion media and filtered with 40 µm Flowmi cell strainers (SP Bel-Art, Wayne, NJ). Cells were transported to the UNM Analytical and Translational Genomics Core for counting and processing. Average processing time from colonoids to library preparation was < 2.5 hours and the cells were highly viable (>90%).

### Single cell and library preparation

Using the Cell Countess II FL (Thermofisher), single cells and viability were counted by Trypan Blue exclusion. The Next GEM Chip G was loaded and put into the Chromium Controller for processing, creating the GEMs. The GEMs were transferred into PCR tubes and cDNA Synthesis was completed per the 10X Chromium Next GEM 3’ Protocol. Synthesized cDNA was evaluated for quality and quantity on a DNA High Sensitivity Chip on the Agilent BioAnalyzer Machine. Next, cDNA was fragmented, adaptor ligated and sample indexed (barcoded) with PCR. Once the libraries were completed, they were processed on a DNA High Sensitivity Chip with the Agilent BioAnalyzer to ensure that the final libraries were approximately 500 bp in size. They were also processed with the KAPA Library Quantification Kit to verify the concentration of final libraries.

### Pooling and sequencing quality control

Sample concentration was determined using Agilent Tape Station 4200 and Invitrogen Qubit 4.0 reagents. Samples were diluted, normalized, and pooled at 4nM prior to qPCR to determine cluster efficiency. Samples were loaded onto Illumina NovaSEQ 6000 instrument and sequenced on S4 Flow cells at specified read length. Data was delivered in FASTQ file format.

### Processing FASTQ reads into gene expression matrices

Raw FASTQ reads from sequencing were processed using Cell Ranger (V2.0.0) software from 10X Genomics. In brief, sequencing reads were demultiplexed, unique molecular identifier (UMI) were collapsed and reads aligned to 10X human transcriptome GRCh38-3.0.0 as the reference.

### scRNA-seq quality control and processing of 10x sequencing data

The raw data created by Cell Ranger was read into the Seurat R package for quality control and downstream processing. Low quality cells expressing <200 genes and genes expressed in fewer than 3 cells were filtered out. After looking at the violin plots of the number of counts, features (genes), and percent mitochondria per sample, we further filtered cells expressing >2000 genes in control samples and cells expressing >7500 genes in UBD treated samples to avoid probable doublets. Cells expressing >20% of mitochondrial genes were filtered out in control and UBD samples. To account for differences in sequencing depth across samples, the data was log-normalized and scaled.

### Clustering, visualization and cell annotation

For cell clustering, we sorted the top 2000 highly variable genes that differed from cell to cell for downstream analysis. The dimensionality of the data was reduced by principal component analysis (PCA) on the highly variable genes. Since Seurat clusters cells based on their PCA scores, the elbow plot was used to determine the top principal components (PCs) for use in clustering the data. From the elbow plot, the top 15 PCs were used in clustering. Cells were divided into unsupervised clusters using a shared-nearest-neighbor modularity optimization-based clustering algorithm. Cells were clustered at a resolution of 0.4. and visualized using Uniform Manifold Approximation and Projection (UMAP) plots (Becht et al., 2018). Differentially expressed genes per cluster were calculated using the Seurat Wilcoxon signed-rank test. Cluster annotation was based off the top genes (log2 fold change > 0.25 and P > 0.05) and marker genes described in literature. Control and UBD treated samples were annotated together to draw direct comparisons. Marker gene expression was visualized using violin plots and dot-plots (size of the dot reflects the percentage of cells expressing the gene and color indicates relative expression). R version 4.0.4 was used for all data plotting and analysis.

### Differences between treatment

For comparisons between control and UBD treated samples, the samples from the same subject were log normalized and scaled separately and were merged into one data set for downstream analysis. Where necessary, all control samples were merged into one dataset and all UBD treated samples were merged into another data set.

### Percentage of cells and statistical analysis

The relative abundance of each cell type was calculated as percentage of cells per condition (control and UBD). We tested for statistical significance using two-tailed t-test (GraphPad Prism 9.0, La Jolla, CA).

### Pathway analysis

Differentially expressed (DE) genes among clusters or samples were ranked according to their expression with the upregulated ones at the top of the list and the downregulated ones at the bottom. The ranked gene list was imported onto gProfiler. Pathway analysis was done using gene ontology (GO) biological process and Reactome. After running the analysis, the gProfiler data was downloaded as a GMT file while the results file was downloaded in the Generic Enrichment Map (GEM) format. The GMT and GEM files were then imported into Cytoscape where the EnrichmentMap app was used to generate the enrichment map. We used a p-value cutoff and q-value cut-off of 1 and an Overlap Coefficient cut-off of 0.5. Using DE genes from Seurat as input, protein-protein interactions of upregulated genes in the UBD treated enteroendocrine cluster were determined using the STRING app on Cytoscape.

### Immunofluorescence, RNAscope and confocal imaging

Colonoids were harvested from Matrigel using Cultrex Organoid Harvesting Solution. They were pelleted (300 xg, 10 min, 4°C), and fixed for 40 min in 4% paraformaldehyde (Electron Microscopy Sciences, Hatfield, PA). Colonoids were permeabilized and blocked simultaneously for 1h using 10% Fetal Bovine Serum (Atlanta Biologicals, Flowery Branch, GA), 0.1% saponin (MilliporeSigma) solution prepared in PBS. After three PBS washes, 100μl of primary antibody (**Supplemental Table 2**), prepared at 1 µg/ml in PBS was added to the cells and incubated overnight at 4°C. Afterwards, cells were washed 3 times with PBS, and 100 μl of AlexaFluor secondary antibodies, Hoechst 33342 (1 mg/ml, all ThermoFisher), and phalloidin iFluor 647 (AAT Bioquest, Pleasanton, CA), (**Supplemental Table 2**), diluted in PBS, were added for 1h at room temperature. After three PBS washes, 50 μl of FluorSave Reagent (MilliporeSigma) was added to the cells and they were mounted between a glass slide and a number 1 coverslip.

Fixed colonoids were prepared for whole mount staining with RNAscope Multiplex Fluorescent V2 using probes Hs-CHGA and Hs-PROX1 (**Supplemental Table 3**). Sample preparation and labeling were performed according to the manufacturer’s protocol (Advanced Cell Diagnostics, Newark, CA). Opal fluorophores (Perkin Elmer, Waltham, MA) and DAPI were used for visualization. 50 μl of FluorSave Reagent was added to the colonoids and they were mounted between a glass slide and a number 1 coverslip. Confocal imaging was carried out in the University of New Mexico Cancer Center Fluorescence Microscopy Shared Resource using an LSM 800 AiryScan confocal microscope (Zeiss) and UNM AIM Center using an LSM 900 AiryScan confocal microscope (Zeiss).

### Statistics

Data are represented as mean ± SEM. Statistical significances were calculated using Student’s *t*-test. Significance was represented as at least p < 0.05. All experiments were performed on a minimum of 3 different colonoid lines derived from separate normal human subjects, with a total of 7 colonoid lines used throughout these studies. N refers to numbers of independent replicates performed. All analyses were performed on GraphPad Prism 9.0.

## Resource Availability

### Lead Contact

Further information and requests for resources and reagents should be directed to the Lead Contact, Julie G. In.

### Materials Availability

All unique reagents generated in this study are available from the Lead Contact without restriction.

### Data and Code Availability

All scRNA-seq datasets were deposited into Gene Expression Omnibus (GEO) and Sequence Read Archive (SRA) and will be released upon publication.

## Author Contributions

MJC, EFC, JGI conceived of experiments. RA, LLA, JGI designed experiments. AB characterized and analyzed uranium-bearing dust composition. RA, LLA, FTL performed the experiments. RA, LLA, FTL, AB, EFC, MJC, JGI analyzed the data. RA, LLA, JGI wrote the manuscript with critical edits from FTL, AB, MJC, EFC. All authors reviewed the final version.

## Acknowledgements

This work was supported by NIH grants K01DK106323 (JGI), R56ES034400 (JGI), P42ES025589 (EFC, JGI), P20GM121176 (EFC, JGI), P20GM130422 (MJC), P50MD15706 (JGI), and American Gastroenterological Association-Aman Armaan Ahmed Family Summer Undergraduate Research Fellowship (LLA). We thank Michael L. Paffett, Jamie L. Padilla, and Kathryn J. Brayer (University of New Mexico Comprehensive Cancer Center, Albuquerque, NM, USA) for technical assistance with confocal microscopy, single cell sequencing, and bioinformatics, respectively. This research was partially supported by the UNM Comprehensive Cancer Center Support Grant NCI P30CA118100.

---

**Supplemental Fig 1.**
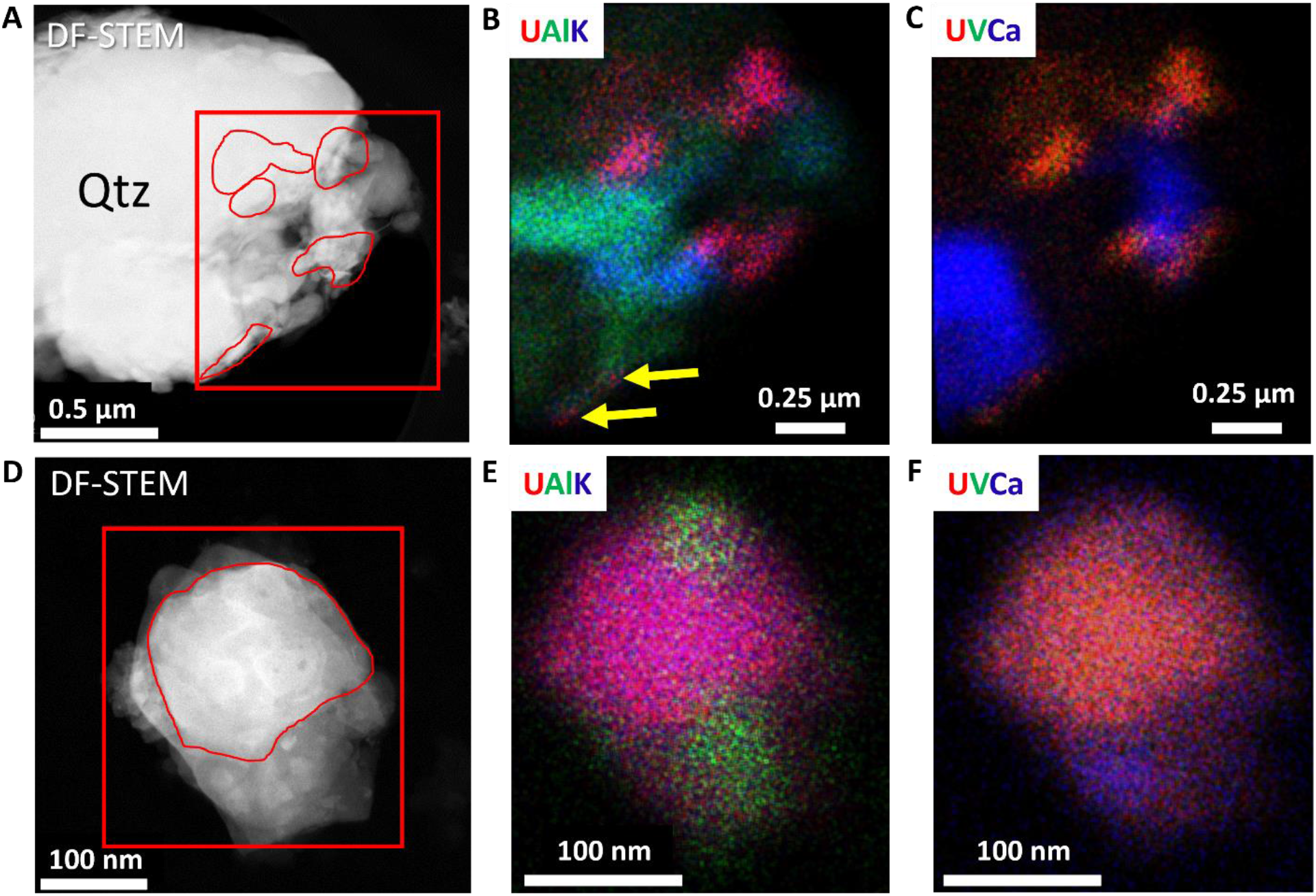
STEM images and STEM EDS X-ray maps of U-bearing particulates. **A)** Dark-field STEM image showing an example of the complex occurrence of U-bearing particulates within the dust sample. A large quartz grain (Qtz) has a region of much smaller grains attached to it on the right-hand side of the image. The outlines of U-bearing grains identified based on STEM EDS analysis are in red. The region of the STEM X-ray map is outlined by the red box. **B)** U M_α_ -Al K_α_ -K K_α_ RGB STEM X-ray map and **C)** U M_α_-V K_α_-Ca K_α_ RGB STEM X-ray map of the boxed region in **A)**. The pink-colored grains in the upper part of the image are U-K-bearing grains <0.3 µM that also contain V, indicated by the orange grains in **C).** Grains of potassium feldspar (blue-green in **B**) are also associated with the carnotite indicating the co-occurrence of U and V in the grains. Two <50 nM carnotite grains are arrowed in **B)**. **D)** Dark-field STEM image of U-bearing particulate associated with silicate minerals. **E)** U M_α_ -Al K_α_ -K K_α_ RGB STEM X-ray map and **F)** U M_α_-V K_α_-Ca K_α_ RGB STEM X-ray map of the boxed region in **D)**.

**Supplemental Fig 2.**
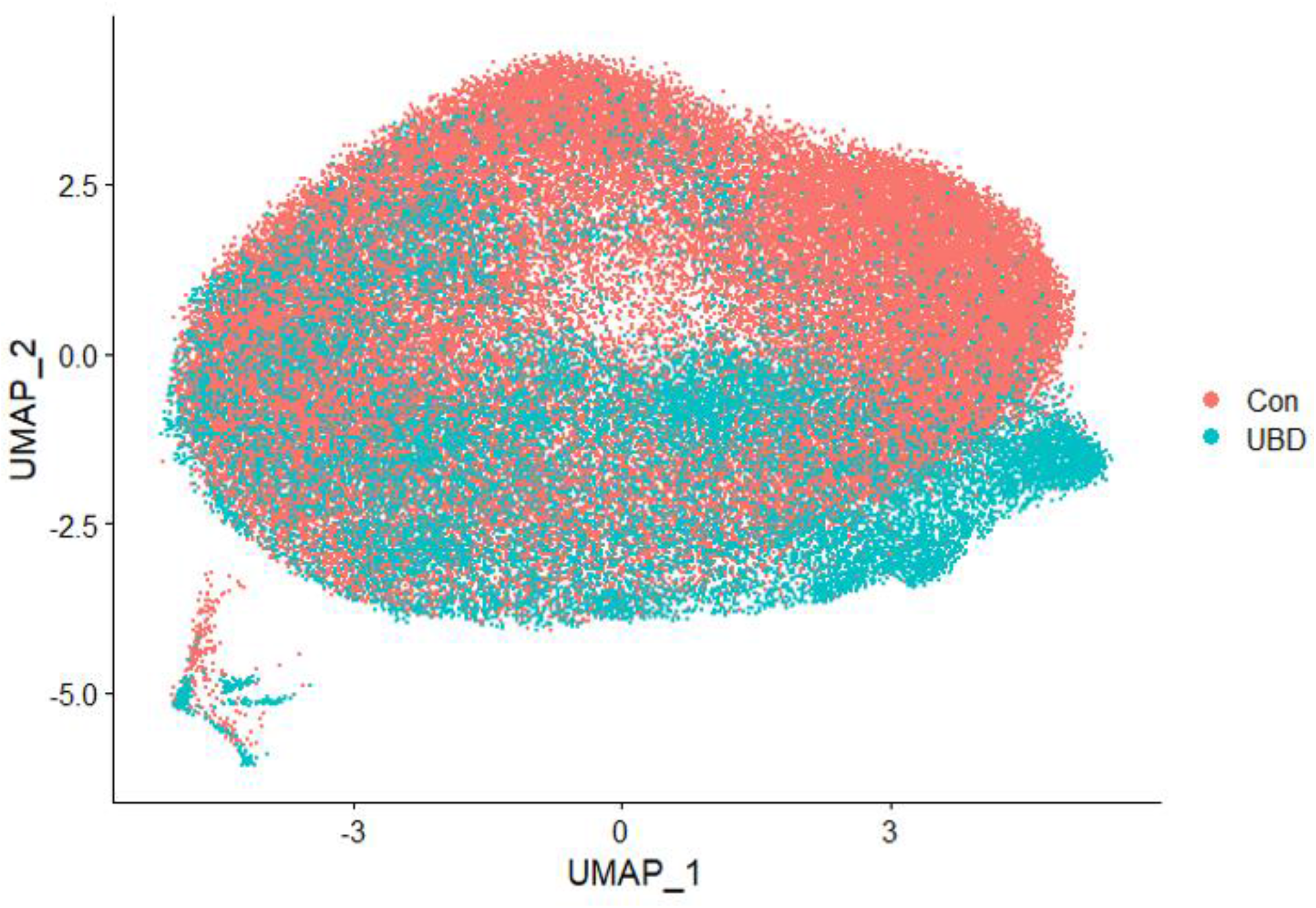
Representative UMAP of merged control and UBD treated colonoids split by treatment. UBD treated cells (blue) do not entirely overlap with control cells (red), indicating changes in gene expression pattern of cells upon UBD exposure.

**Supplemental Fig 3.**
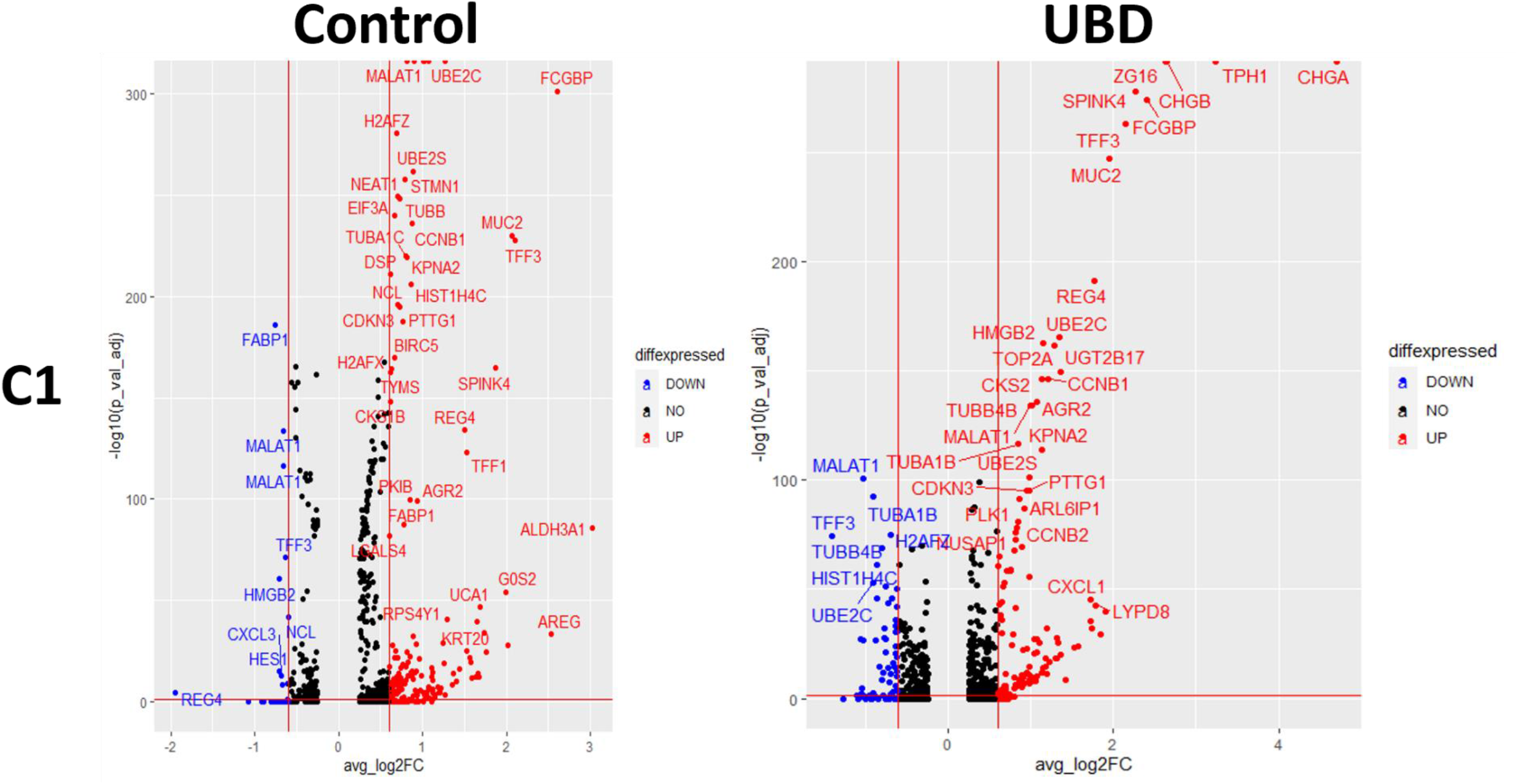
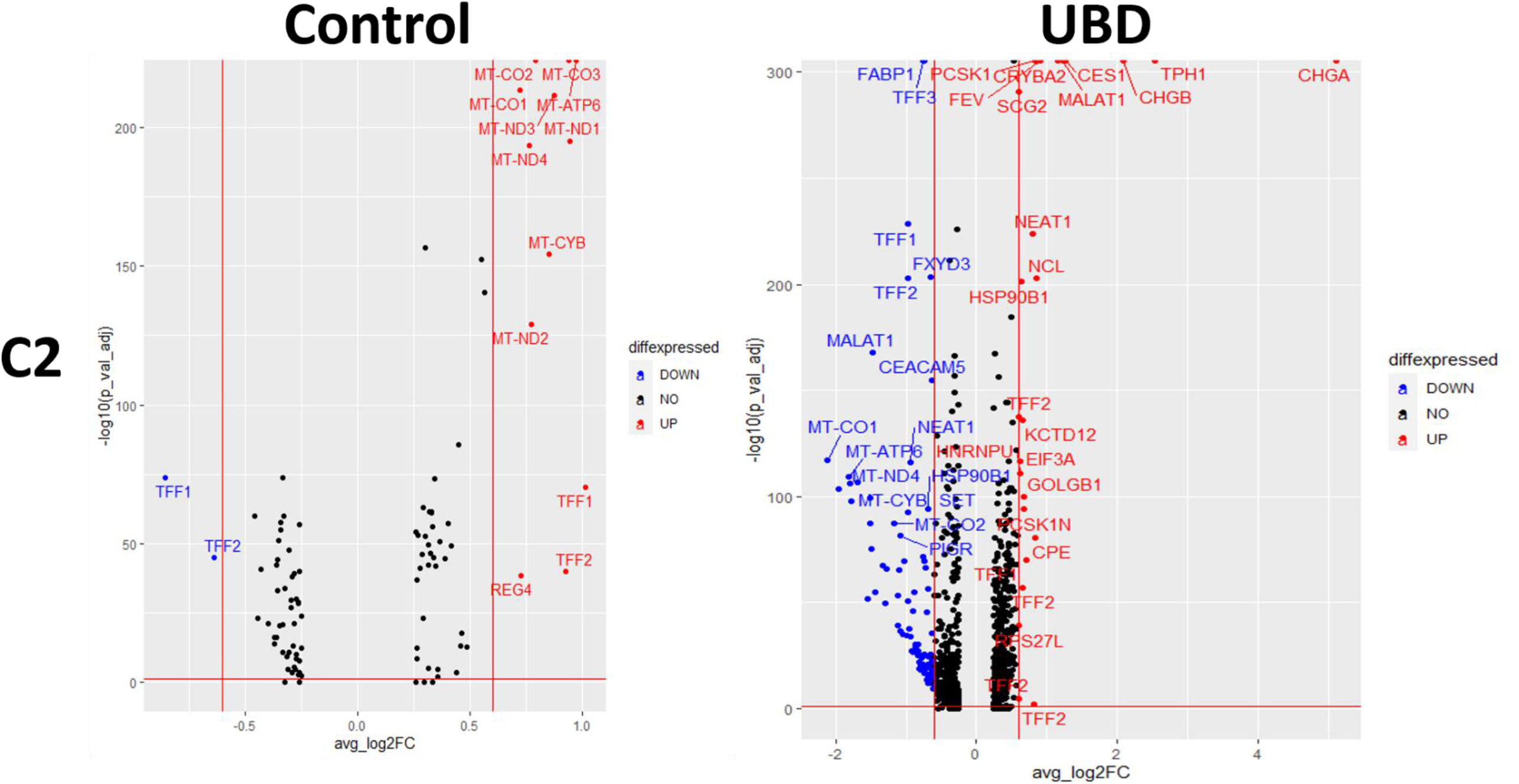
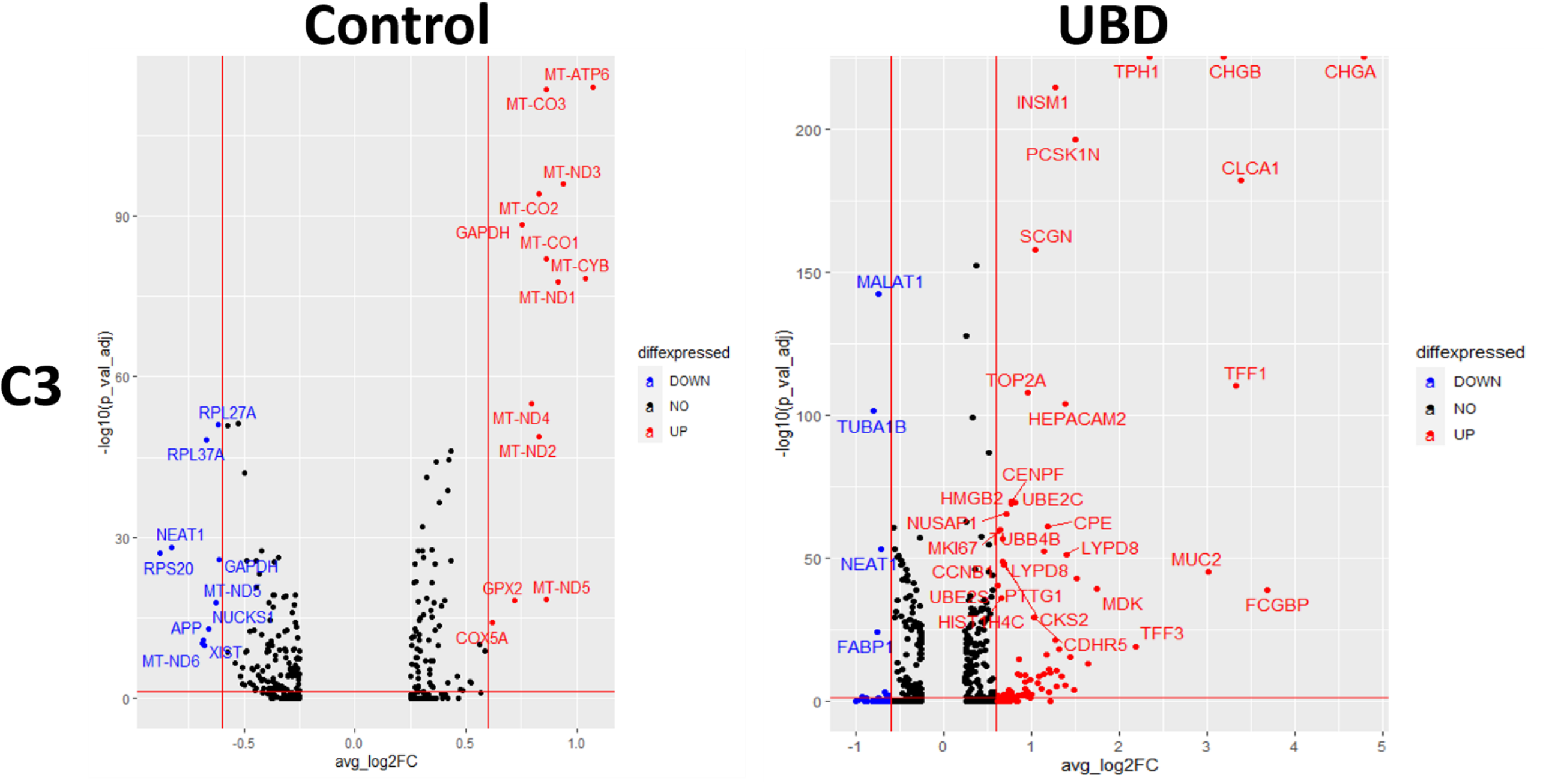
Volcano plots of control and UBD treated colonoids split by treatment. Significantly up (red) and down (blue) regulated genes in all three sequenced colonoid lines were separated by treatment. Genes associated with secretory lineage (goblet, enteroendocrine) were upregulated in all UBD treated colonoid lines.

**Supplemental Fig 4.**
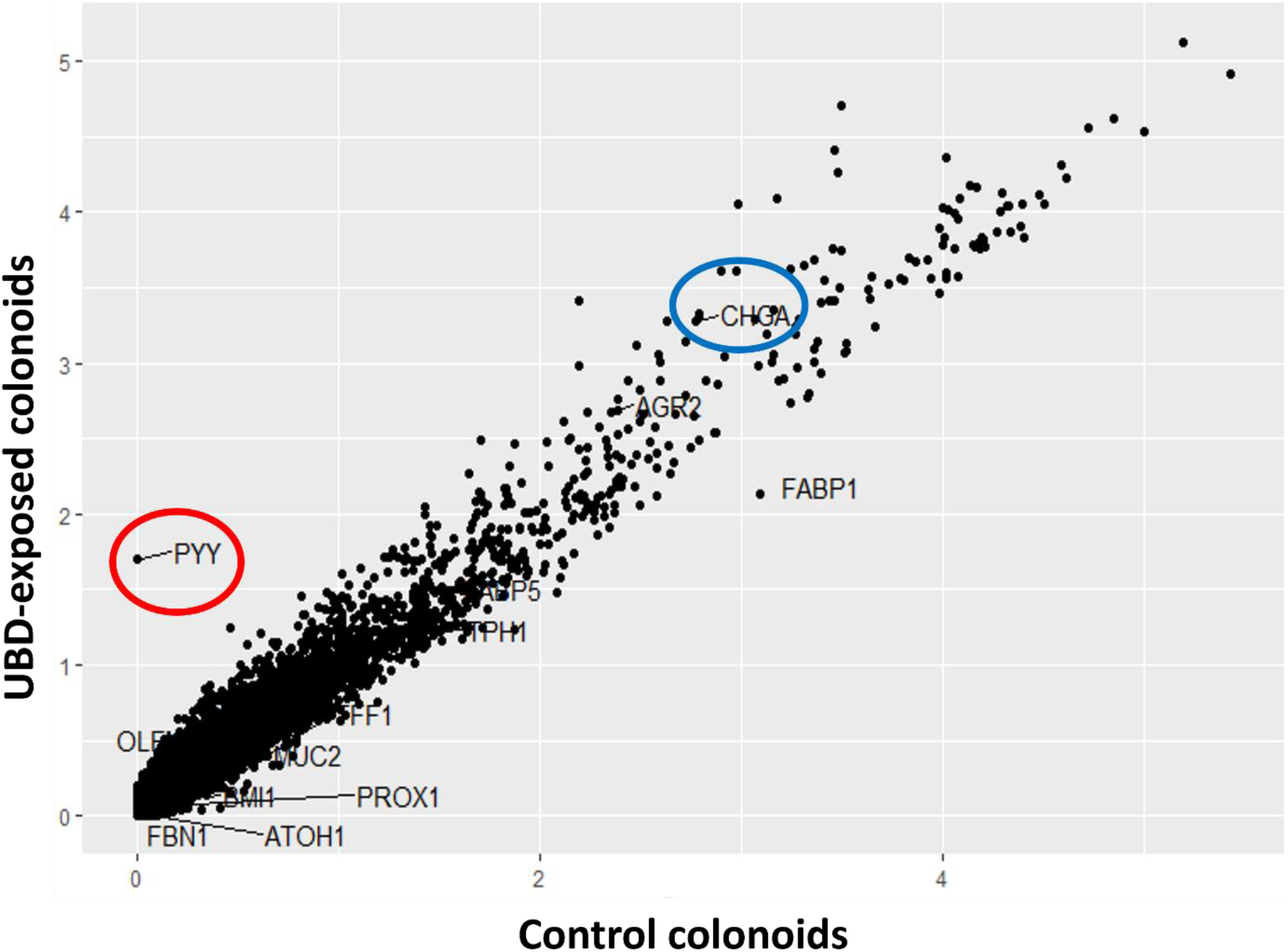
Scatter plot of average gene expression in EECs by treatment. PYY expression (L cells, circled in red) is nearly absent in EECs in control colonoids. CHGA (total EECs, circled in blue) is higher in UBD treated EECs.

**Supplemental Table 1.** Description of colonoid donors used for scRNA-seq. All colonoids were generated from proximal colonic biopsies from patients undergoing routine, screening colonoscopy and with no pathophysiology.

| **Colonoid identifier**<br>**(scRNA-seq)** | **Age** | **Sex** | **Race** |
| --- | --- | --- | --- |
| **C1** | **70** | **M** | **Hispanic, non-white** |
| **C2** | **58** | **M** | **White** |
| **C3** | **67** | **F** | **African-American** |

**Supplemental Table 2.** Antibodies used for immunostaining. Source and identifier of each antibody used is detailed in the table.

| **Antibody** | **Source** | **Identifier** |
| --- | --- | --- |
| 5-HT | Fitzgerald | 5-HT-H209 |
| PYY | Abcam | Ab131246 |
| CHGA | DSHB | CPTC-CHGA-1 |
| CHGA | Santa Cruz | SC393941 |
| PROX1 | Novus | NBP1-30045 |
| Donkey anti-Rabbit IgG-AF488 | ThermoFisher | A-21206 |
| Donkey anti-Mouse IgG, AF568 | ThermoFisher | A10037 |
| Donkey anti-Mouse IgG, AF488 | ThermoFisher | A21202 |
| Goat anti-Rabbit IgG AF568 | ThermoFisher | A1101042 |
| Phalloidin-ifluor 647 | AAT Bioquest | 1000854 |

**Supplemental Table 3.** RNA oligo probes used for RNAscope. Probes were all sourced from ACD. Identifiers for each probe is detailed in the table.

| **Probe** | **Source** | **Identifier** |
| --- | --- | --- |
| Hs-CHGA-C4 | Advanced Cell<br>Diagnostics | 311111-C4 |
| Hs-PROX1-C2 | Advanced Cell<br>Diagnostics | 530241-C2 |

